# Centrosome positioning independently of microtubule-based forces

**DOI:** 10.1101/2024.06.11.598451

**Authors:** Alexandre Schaeffer, Morgan Gazzola, Matthieu Gelin, Benoit Vianay, Chiara de Pascalis, Laurent Blanchoin, Manuel Théry

## Abstract

The regulation of centrosome position is central to the establishment of polarized cell functions. This process is thought to rely on the generation of mechanical forces along microtubules, the balance of which determines centrosome position. By studying these forces in adherent cells in culture, we found that neither pushing nor pulling forces propagate along microtubules to the centrosome. Inhibiting dyneins or disassembling microtubules did not disrupt the maintenance of the centrosome at the center of the cell. In contrast, the actomyosin contractile network appears to be responsible for the generation of a centripetal flow that drives the centrosome towards the center of the cell independently of the microtubules. Furthermore, we found that centering of the centrosome depends more on the reorganization of cell shape around the centrosome than on an effective centrosome displacement throughout the cytoplasm. Interestingly, despite their lack of mechanical role, microtubules appear to direct this remodeling of cell shape. This revised view of centrosome positioning offers a new perspective for understanding the establishment of cell polarity.

## Introduction

In animal cells, the centrosome is the main microtubule-organizing center. By nucleating and anchoring microtubules, it organizes the network in a polarized radial array, with microtubule minus-ends in and plus-ends out. The centrosome is therefore responsible for the orientation and direction of intra-cellular traffic and the organization of cell polarity. Observing the centrosome in early sea urchin and Ascaris embryos, Theodore Bovery named it for its position at the center of the cell’s soma (Scheer, 2014). The name stuck because this remarkable geometric property was quite robust and could be easily observed in most adherent cells in culture. Interestingly, in many conditions, such as polarized epithelial cells or immune cells, the centrosome is located at the periphery of the cell. This suggests that the central positioning of this organelle is not a fixed rule and that specific mechanisms may regulate it differently depending on the cellular context (Tang and Marshall, 2012). A better understanding of this process is essential as the regulation of centrosome positioning is involved in many important physiological functions ranging from directed intracellular transport, cell locomotion, immunological response as well as cell proliferation and differentiation. (Hannaford and Rusan, 2024). However, numerous studies carried out in different organisms and cell types have revealed distinct and apparently contradictory characteristics, particularly with regard to the origin and orientation of the mechanical forces acting on the centrosome.

The mechanism of centrosome positioning has been best characterized in eggs and early embryos (Haupt and Minc, 2018). The large size of these cells has enabled direct manipulations and precise mapping of the mechanical forces acting along the microtubule network. Centering appears to rely on both pushing forces produced by the polymerization of microtubules against the cell periphery (Garzon-Coral et al., 2016; Sulerud et al., 2020; Meaders et al., 2020) and on the pulling forces generated by dyneins as they walk toward microtubule minus-ends (Grill et al., 2003; Tanimoto et al., 2016; Farhadifar et al., 2020; Wu et al., 2024). Both mechanisms can ensure a robust centering of the centrosome (Kimura and Kimura, 2011; Grill and Hyman, 2005; Pecreaux et al., 2016). However, the large scale coordinated motions of microtubule asters, organelles and actin filaments in Xenopus egg extracts is not compatible with the application of forces along microtubules only and rather suggested that all components of the cytoplasm behave as an physically integrated gel (Pelletier et al., 2020).

In somatic cells, and particularly in adherent cells in culture, the positioning mechanism is much less clear. Physical forces have not been directly assessed and the parameters involved in centering have been deduced from the cell’s response to biochemical changes or geometric constraints. Although dynein-based pulling forces along microtubules have been argued to participate to centrosome centration, dynein inactivation or microtubule disassembly did not completely randomize centrosome position (Burakov et al., 2003; Wu et al., 2011; Hale et al., 2011). This could be explained by the additional contribution of the actin network, which has been proposed to push directly on microtubules (Burakov et al., 2003; Hale et al., 2011) or simply to provide physical boundaries to the microtubule network (Jimenez et al., 2021). In addition, and in contrast to the radial arrangement of straight microtubules in eggs, the irregular organization of many microtubules and their tortuous shapes in adherent cells in culture seem incompatible with most models based on either pushing or pulling forces along them (Zhu et al., 2010; Letort et al., 2016). Thus, the origin and spatial distributions of mechanical forces acting on the centrosome in somatic cells were unclear and required further investigation.

## Results

To directly reveal the exact contributions of pushing and pulling forces to the mechanical force balance defining the position of the centrosome, we used laser-based nano-ablation to cut microtubules. We worked with PtK2 cells stably expressing tubulin-GFP in order to visualize the microtubules. Unexpectedly, ablation of a few microtubules near the centrosome had no significant impact on centrosome position (Figure 1A and movie S1). Considering that the attachment of the centrosome to the nucleus might buffer small changes in the forces acting on it (Salpingidou et al., 2007; Lombardi et al., 2011), we repeated the experiment in cytoplasts, ie enucleated cells, but observed no further effect (Figure 1B and movie S2). To challenge the model of centering based on the balance of mechanical forces, we then ablated about half of the microtubules reaching the centrosome. As the loss of dynamic microtubules can be rapidly compensated by the growth of new microtubules, we continuously ablated all microtubules reappearing in the ablated regions over the next few minutes (Figure S1A, B). However, this harsh surgery only led to a barely visible slow recoil (less than half a micron in 3 to 5 minutes) of the centrosome away from the ablated region (Figure 1C and movie S3). These data called into question all previous models of force balance at the centrosome, and we wondered whether they might result from the specific shape and intracellular organization of PtK2 cells. We therefore repeated the experiment in other cell types with clearly distinct cytoskeleton architectures: RPE1 cells with smaller sizes and denser microtubule arrays, MEF cells with more irregular shapes and far fewer microtubules (Figure S1C). In cytoplasts from both cell types, centrosome recoil after microtubule ablation was as small and slow as in PtK2 cells (Figure S1D), and markedly different from the 5 micrometers displacement in about 20 seconds that had been observed following microtubule ablation in early *Caenorhabditis elegans* embryos (Farhadifar et al., 2020; Wu et al., 2024). Interestingly, in all cell types, and in the few cases of clear centrosome recoil, a similar relaxation was also visible in the surrounding actin network (Figure 1D, S1E,F). This co-relaxation suggested that the two networks were bound to each other. It also suggested that the centrosome displacements we observed were actually due to the severing of actin filaments bundles by the ablation rather than specific severing of microtubules.

**FIGURE 1:**
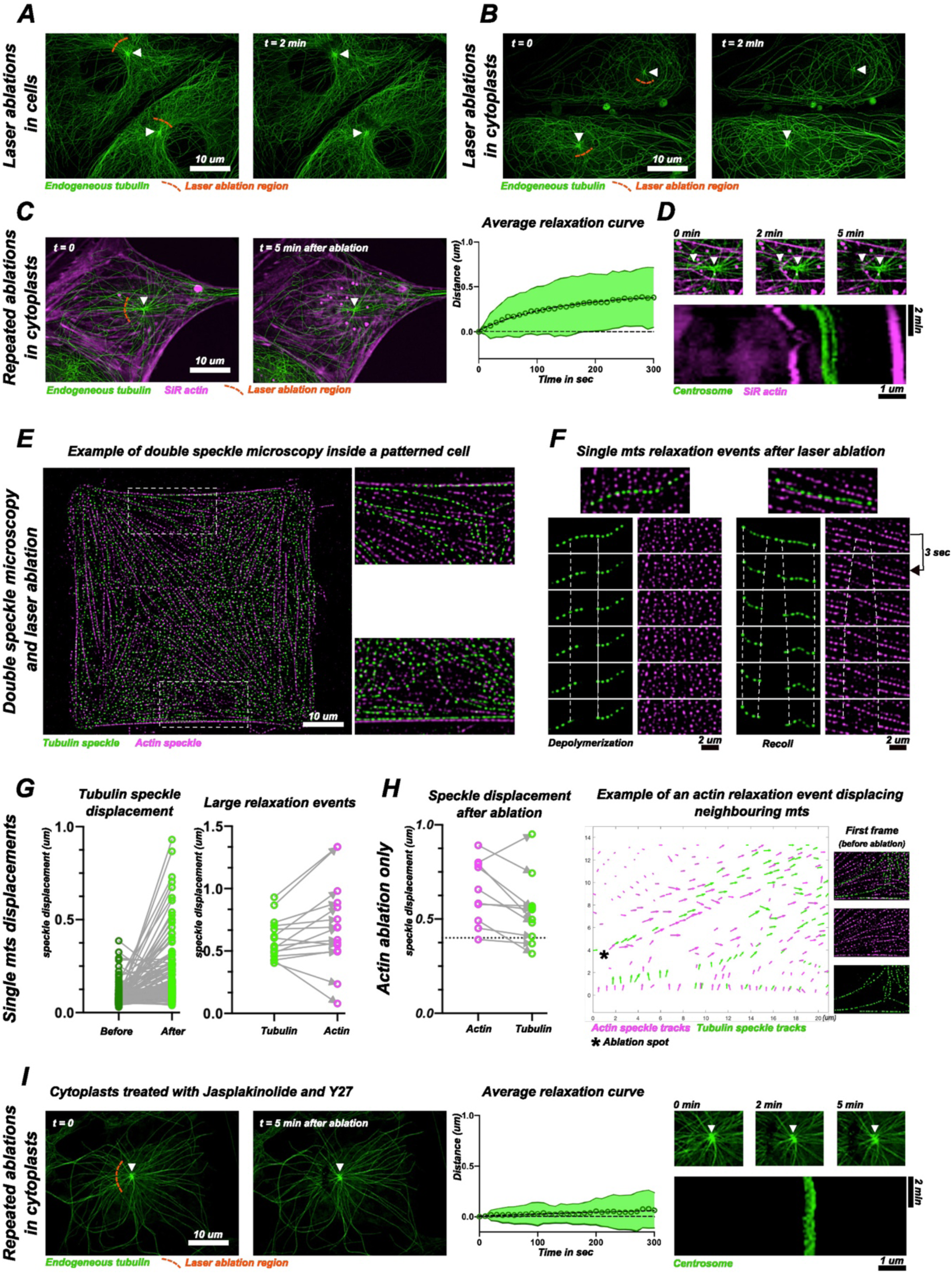
Relaxation of microtubules upon laser ablation. ***(A)*** First and last timepoints of a 2 minutes laser ablation experiment in PtK2-GFP cells. A few microtubules were ablated on one side of the centrosome. White arrowheads indicate the positions of the centrosomes. ***(B)*** First and last timepoints of a 2 minutes laser ablation experiment in PtK2-GFP cytoplasts. A few microtubules were ablated on one side of the centrosome. White arrowheads indicate the positions of the centrosomes. ***(C)*** First and last timepoints of a laser ablation experiment in a PtK2 cytoplast (left). Actin was stained using SiR Actin. Microtubules were extensively and repeatedly ablated on one side of the centrosome during five minutes. White arrowheads indicate the positions of the centrosomes. Graph displays the average relaxation curve showing the mean relaxation profile of the centrosomes (n=22) during the 5 minutes laser ablation experiment. The displacement of the centrosome was projected along an axis connecting the centroid of the ablated area and the centrosome (for a graphical representation see supplementary figure 1A). The circles represent the average displacement of the centrosome at each timepoint, the continuous lines represent the standard deviation. ***(D)*** Zoomed in view of the centrosomal area of the cytoplast shown in figure 1C. White arrows indicate from left to right a recoiling actin structure and the centrosome position. Kymograph shows centrosome and actin relaxations in the zoomed area. The linescan was performed along a line connecting the centrosome with the ablated area and spanning the entire length of the zoomed in area. ***(E)*** Representative live image of a PtK2 cell platted on a 3500um^2^ H pattern and microinjected with both actin and tubulin. On the right, two magnified regions showing in higher details actin and tubulin speckles. ***(F)*** Representative responses of single microtubules in the 15 seconds following laser ablation. Left images show microtubule depolymerization after laser ablation with no mechanical relaxation. Right images show local microtubule buckling and recoiling and actin stress fiber recoiling after laser ablation. ***(G)*** Graphs show the displacements of tubulin speckles before and after ablation (n=121, left), and the displacements of tubulin and actin speckles after laser ablation in the cases of large relaxation events (displacements > 400nm) (n=15, right). ***(H)*** Microtubule displacement events occurring when only actin was ablated. Graph shows the motion of tubulin speckle and actin speckle after ablation. The dotted line marks the 400 nm threshold used to define large relaxation events (n=10). Vector map shows the displacements of actin and tubulin speckles upon ablation of actin bundle (over a 15 second period). Images show actin and tubulin speckles localization before the laser ablation event. ***(I)*** Images on the left show the first and last timepoints of a laser ablation experiment in a PtK2-GFP cytoplast treated with Jasplakinolide (600nm) and Y27632 (20µM) for 4 hours. Microtubules were extensively and repeatedly depleted on one side of the centrosome during five minutes. White arrowheads indicate the positions of the centrosomes. Graph displays the average relaxation curve showing the mean relaxation profile of centrosomes (n=24) during the 5 minutes laser ablation experiment. The circles represent the average displacement of the centrosome at each timepoint, and the continuous lines represent the standard deviation. Images on the right show a magnified view of the centrosomal area of the cytoplast shown in figure 1I. White arrowheads indicate the centrosome position. Kymograph shows centrosome relaxation in the magnified area. The linescan (scaled 3 times to smoothen the signal) was performed along a straight line connecting the centrosome with the centroid of the ablated area and spanning the entire length of the zoomed in area. ***(A B C I)*** All images are max projections, further processed using an unsharp mask, a gamma filter and a subtract background function. ***(E F)*** *For* the details regarding actin and tubulin speckle processing, see the dedicated sections in the extended material and methods.

We decided to investigate further the possible entanglement of the two networks and analyzed the consequences of severing one of them on the relaxation of the other. We micro-injected low doses of labelled tubulin and actin in order to generate speckles along the two networks and precisely monitor filament displacements (Salmon et al., 2002). The tubulin speckles allowed us to distinguish microtubules polymerization, depolymerization or pause from translocation events themselves (Figure S2A and movie S4). Similarly, actin speckles revealed filament translocation along stress fiber and fiber displacements (Figure S2B and movie S5). Cells were plated on adhesive micropatterns to normalize the architecture of the actin network. We used geometries were cells could spread over non adhesive regions to induce the formation of contractile bundles that would relax more freely (Vignaud et al., 2021) (Figure 1E, and Figure S2C). The recoil of actin bundles and microtubules was measured by tracking individual speckles displacements (Figure S2D). Local ablations in those regions led to either microtubule disassembly with no detectable effect on actin bundles, or to the recoil of both microtubules and actin bundles (Figure 1F and movie S6). Most microtubule recoils were shorter than 200 nm. The few recoils that were larger than 400 nm were associated with a similar recoil of the local actin network which was also disrupted during laser ablation (Figure 1G and Figure S2E). Conversely, ablation of actin bundles was consistently associated with microtubule recoil. Importantly, specific severing of actin bundles in regions devoid of microtubules also resulted in the displacement of distant microtubules (Figure 1H and movie S7). These experiments therefore showed that microtubules were not free in the cytoplasm but rather attached to actin bundles along their length. This was confirmed by the absence of centrosome relaxation following large and repeated ablations of microtubules in conditions where the actin network was “frozen” by blocking actin filament dynamics (with Jasplakinolide) and translocations (with ROCK inhibitor) (Peng et al., 2011) (Figure 1I). These results further challenged the possibility that forces could be generated along microtubules and propagated to the centrosome. They prompted us to re-examine the role of dyneins, which have been shown to be major regulators of centrosome positioning through their production of pulling forces along microtubules in multiple systems (Hooikaas et al., 2020; Koonce, 1999; Tanimoto et al., 2016; Nguyen-Ngoc et al., 2007; Farhadifar et al., 2020; Wu et al., 2024).

We tested the effect of two potent and specific dynein inhibitors: Dynapyrazole and Dynarrestin. Dynapyrazole blocks dynein’s ATPase activity (with an IC50 10 µM) (Steinman et al., 2017) and Dynarrestin blocks dynein’s binding to microtubules (with an IC50 5µM) without interfering with the ATPase activity (Höing et al., 2018). Treating PtK2 cells for 1 hour with either Dynarrestin (50 µM) or Dynapyrazole (80 µM) induced a fragmentation of the Golgi apparatus into multiple small stacks (Figure S3A, B), but did not result in their dispersion throughout the cell, as would occur with complete dynein inactivation (Harada et al., 1998) (Figure 2A, B). Interestingly, the distance between the centrosome and the center of mass of the cell was not affected (Figure 2C). We then induced a stronger inactivation of dyneins by injecting a blocking antibody against dynein (Figure 2D) (Dillman and Pfister, 1994). In contrast to dynein inhibitors, this led to a clear dispersion of the Golgi apparatus (Figure 2E) (Yi et al., 2011). However, dynein blocking antibodies did not affect the positioning of the centrosome either (Figure 2F), even though we could observe an important perturbation of the pericentrosomal microtubule network with a loss of its astral organization (Figure 2G) (Quintyne et al., 1999). The various dynein inhibition strategies can either interfere with the ATPase activity of dyneins or with their ability to bind to microtubules or to their cargoes. They were thus reported to have various and sometimes opposite effects on Golgi fragmentation and dispersal (Quintyne et al., 1999; Barr and Short, 2003; Yi et al., 2011; Jimenez et al., 2021) and on the centering and reorientation of centrosomes (Palazzo et al., 2001; Burakov et al., 2003; Yi et al., 2013) in distinct cell types and in distinct conditions. To further strengthen our conclusions on PtK2, we also tested these inhibitions strategies on MEF and RPE1 cells, which have different cytoskeleton architectures. In both cell types, as in PtK2, the Golgi apparatus was dispersed by the injection of blocking antibodies (Figure S3E, I) but not by the drugs (Figure S3C, G). The radial organization of microtubules was perturbed in MEF but not in RPE1 cells (Figure S3F, J). In both cell types, as in PtK2 cells, the centrosome remained close to the cell center in response to all dynein inhibition strategies (Figure S3D, H). These results showed that the centering of the centrosome in adherent cells in culture does not require the binding or the activity of dyneins. Counter-intuitively, a strong perturbation of the architecture of the pericentrosomal microtubule network did not affect the positioning of the centrosome. In light of our previous results on microtubule mechanics, this further questioned the actual role of microtubules in centrosome positioning.

**FIGURE 2:**
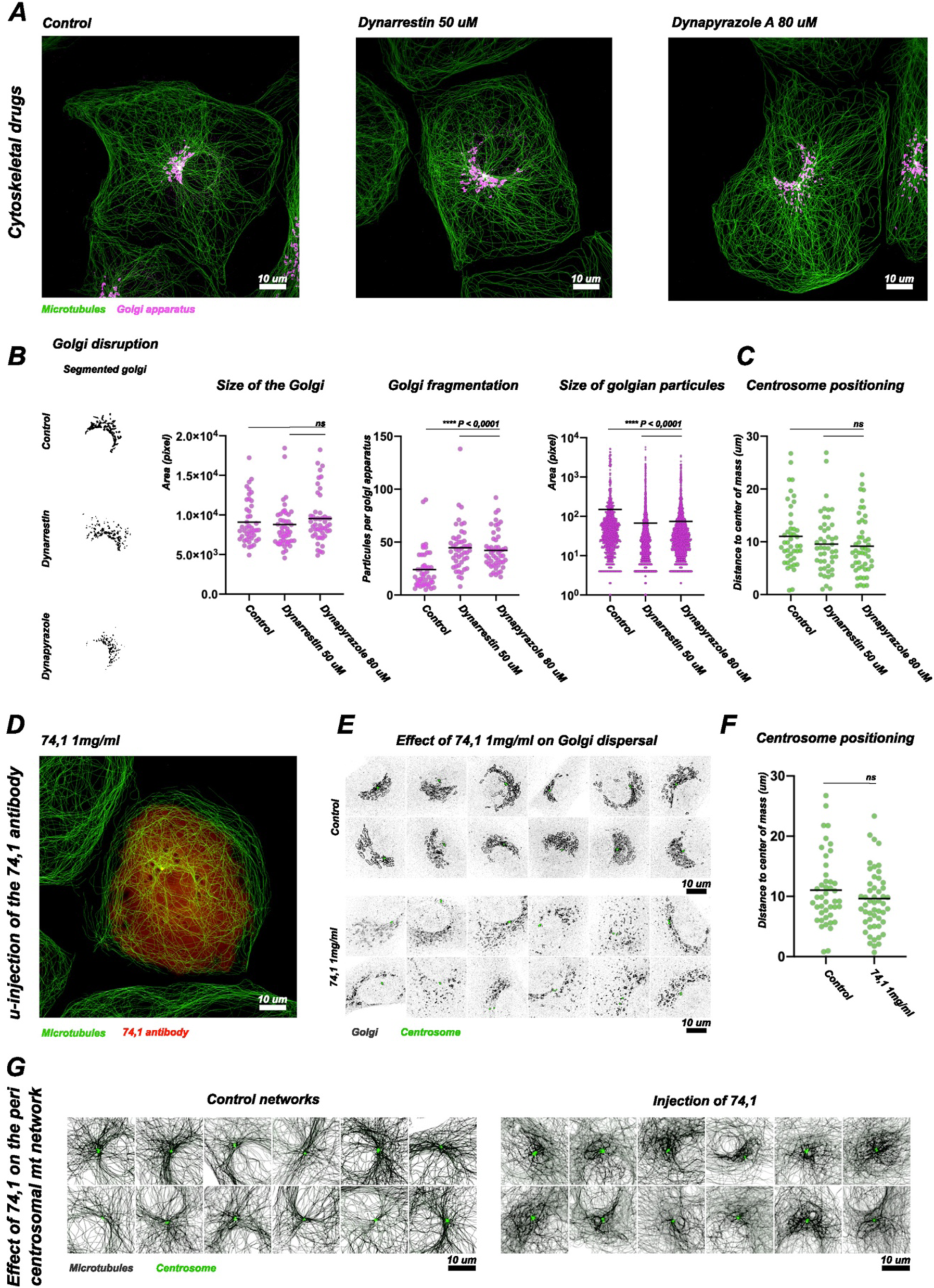
Effect of dynein inhibition on centrosome positioning. The activity of dynein molecular motors was inhibited inside PtK2-GFP using a one-hour treatment of Dynarrestin (50µM), Dynapyrazole A (80µM) or through the micro-injection of the 74.1 (1mg/ml) inhibiting antibody. After micro-injection cells underwent a 50-minute incubation period. In all conditions cells were fixed and stained for the microtubules, the centrosome, the Golgi apparatus, and actin. For each inhibitor the extent of dynein inhibition is assessed using the morphology of the Golgi apparatus. In the graphs, horizontal bars represent the mean. P represents the p values, which were obtained from Kruskal-Wallis non-parametric tests. For each condition, at least 40 different cells or cytoplasts were analyzed. ***(A)*** Representative images of fixed PtK2 cells in control condition (left), one hour of dynein inhibition using Dynarrestin (middle), and one hour of dynein inhibition using Dynapyrazole A (right). ***(B)*** Effect of a 1-hour treatment of Dynarrestin or Dynapyrazole A on the morphology of the Golgi apparatus. Images show the segmented Golgi apparatus in the control and dynein inhibited conditions. Histograms show the size of the Golgi apparatuses in the various dynein inhibited condition, the number of segmented particles in every segmented Golgi apparatus in the control and dynein inhibited conditions, and the size of segmented particles inside every analyzed Golgi apparatus in the control and dynein inhibited conditions (control (n=44), Dynarrestin (n=47) and Dynapyrazole A (n=49) conditions) (for more details see supplementary figure 3A and the dedicated section in the extended material and methods). ***(C)*** Histograms show the distance between the centrosome (whose position was determined using the gamma tubulin staining) and the center of mass of the cell (whose periphery was determined using the actin staining) in PtK2 cells, in the control (n=44), Dynarrestin (n=47) and Dynapyrazole A (n=49) conditions. ***(D)*** Representative image of fixed a PtK2 cell 50 minutes after its micro-injection with the 74.1 (1mg/ml) antibody. ***(E)*** Effect of the injection of the 74,1 (1mg/ml) antibody on the morphology of the Golgi apparatus. Images show Golgi apparatuses in fixed RPE1 cells in control condition (top), and 50 minutes after their microinjection with the 74,1 antibody (bottom). ***(F)*** Histograms show the distance between the centrosome (whose position was determined using the gamma tubulin staining) and the center of mass of the cell (whose periphery was determined using the actin staining) in the control (n=44) and the 74,1 (n=52) micro-injected conditions. ***(G)*** Effect of the injection of 74,1 on the pericentrosomal microtubule network in PtK2 cells. Images show the pericentrosomal network of microtubules in control cells (left), and 50 minutes after the micro-injection with the 74.1 (1mg/ml) antibody (right).

Microtubules are the nexus of numerous signaling pathways (Janke and Magiera, 2020). It is therefore challenging to test their direct contribution to centrosome positioning without impacting the various processes they regulate, which could also be involved in centrosome positioning (Seetharaman and Etienne-Manneville, 2020). In particular, microtubules regulate acto-myosin contractility by sequestering the Rho GEF GEF-H1 (Krendel et al., 2002).The consequence of their disassembly following nocodazole treatment may be confounded with the associated increase of acto-myosin contractility (Rafiq et al., 2019; Seetharaman et al., 2021). To limit this effect, we co-treated cells with nocodazole and an inhibitor of Rho kinase. At low dose (1 µM), Y27632 was not able to fully compensate for the over-contraction associated to the disassembly of microtubules induced by 6µM of Nocodazole. On the other hand, a high dose (30 µM) of Y27632 led to the disassembly of most actin contractile structures in cells treated with 6µM of Nocodazole (Figure 3A). In both cases, most microtubules were depolymerized and only a few short microtubules remained at the centrosome (Figure 3A and Figure S4A-D). Although these few microtubules could not interact with the cell periphery, nor with most of the intracellular space, the centrosome was still properly located at the cell’s center of mass (Figure 3B). As this could result from the anchoring of the centrosome to the cell nucleus, we repeated these treatments in cytoplasts and confirmed that the “mini-asters” remained well centered, without contacting the periphery of the cell and in the absence of any interactions with the nucleus (Figure 3C and D). These results demonstrated that microtubules were not required to maintain the centrosome at the center of the cell. This led us to hypothesize that another cytoplasmic structure might be responsible for the integration of the entire cell volume in the centering process. To challenge the role of the actin network in this process, we treated cells with 0,5 µg/mL of Cytochalasin D to block the assembly of actin filaments and disrupt most actin-based structures. This treatment could not be applied to cytoplast without inducing their detachment, but a significant proportion of cells could remain at least partially spread. After 6h of treatment with Cytochalasin-D the centrosome positions were largely randomized (Figure 3E, F). The dispersion relative to cell size was even more striking (Figure 3G), since these cells were smaller than control cells (Figure S4E). In response to actin filament disassembly, a few centrosomes were even observed at the periphery of the cells (Figure 3E and Figure S3C). Overall, the actin network, and not the microtubule network, appeared to be necessary to maintain the centrosome at the center of adherent cells.

**FIGURE 3:**
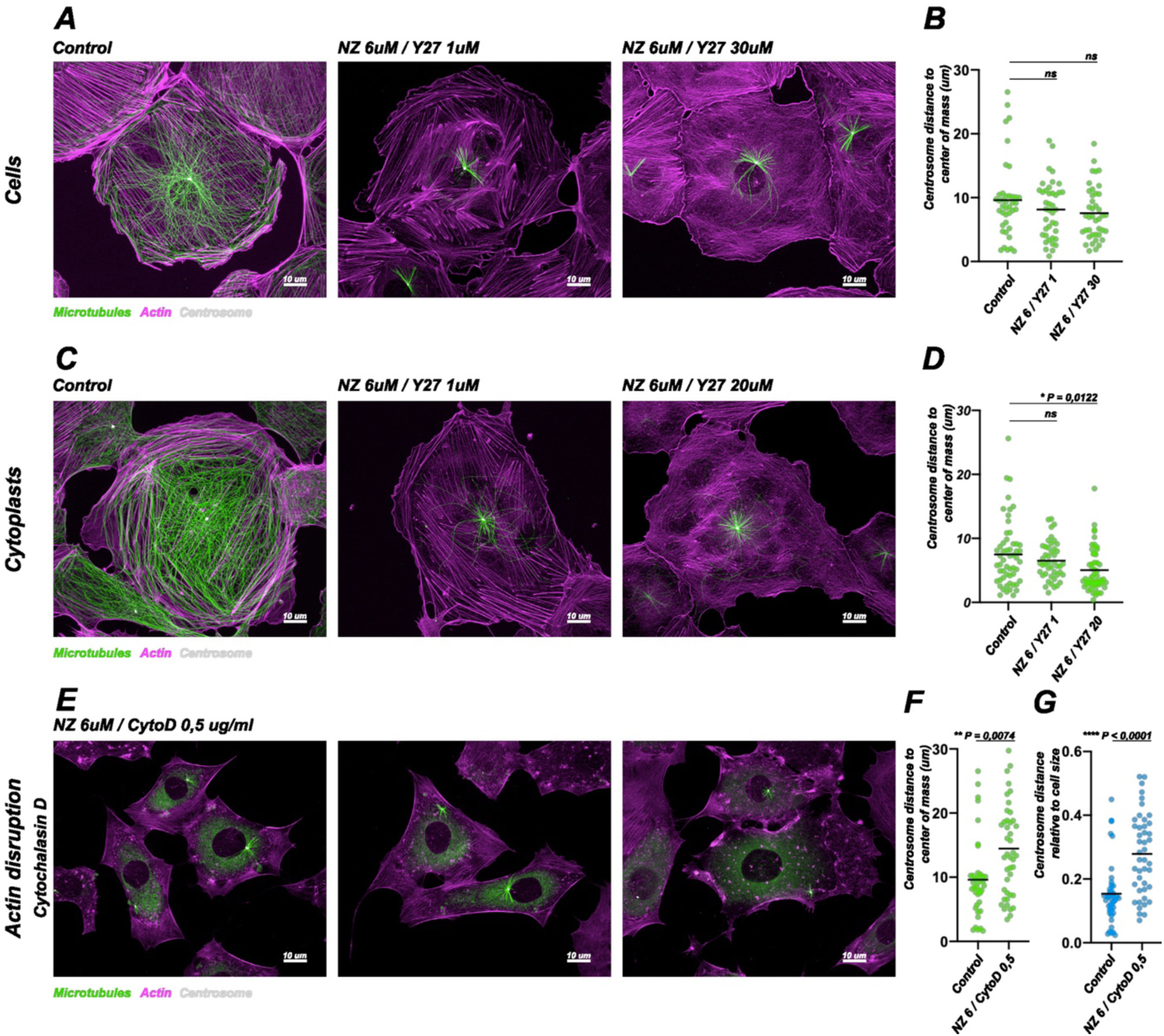
Respective roles of actin and microtubule networks in centrosome positioning. Mini-aster positioning in PtK2-GFP cells ***(A, B, E, F, G)*** and cytoplasts ***(C, D)***. Briefly, cells or cytoplasts were exposed to 6 hours of Nocodazole and Y27632 or Cytochalasin-D at various doses to generate small microtubule asters in strongly contractile, poorly contractile, or disrupted actin networks. The cells and cytoplasts were then fixed and stained for actin, tubulin and the centrosome. In the graphs, horizontal bars represent the mean. P represents the p values, which were obtained from Kruskal-Wallis non-parametric tests. For each condition, at least 40 different cells or cytoplasts were analyzed. ***(A)*** Images show the actin (magenta) and microtubule (green) networks in a control PtK2 cell (left), a cell treated with 6µM of Nocodazole 6µM and 1µM of Y27632 (middle) or 30 µM of Y27632 (right). ***(B)*** Histograms show the distance between the centrosome (whose position was determined using the gamma tubulin staining) and the center of mass of the cell (whose periphery was determined using the actin staining) in the control (n=40), the highly (n=40) and poorly (n=41) contractile cells. ***(C)*** Images show the actin (magenta) and microtubule (green) networks in a control PtK2 cytoplast (left), a cytoplast treated with 6µM of Nocodazole 6µM and 1µM of Y27632 (middle) or 20 µM of Y27632 (right). ***(D)*** Histograms show the distance between the centrosome and the center of mass of the cytoplast in the control (n=55), the highly (n=45) and poorly (n=51) contractile cytoplast conditions. ***(E)*** Images show the actin (magenta) and microtubule (green) networks in a PtK2 cell treated with Nocodazole (6µM) and Cytochalasin-D (0,5ug/mL). ***(F)*** Histogram shows the distance between the centrosome and the center of mass of the cell in the control (n=40) and Cytochalasin D treated cells (n=44). ***(G)*** Histogram shows the centrosome distance to the center of mass normalized by cell size (square root of the area of the cell) in the control (n=40) and Cytochalasin D treated cells (n=44). ***(A C E)*** All images are max projections, further processed using an unsharp mask and a gamma filter.

These results seemed to contradict previous studies concluding that microtubules play a role in the generation and transmission of forces that regulate aster centering in adherent cells (Wu et al., 2011; Hale et al., 2011; Zhu et al., 2010). This includes our own observation of centrosome mispositioning in response to dynein inactivation by overexpression of a dominant negative form of the p150 subunit of dynactin (Jimenez et al., 2021). We wondered whether the long-term effects of protein knockdown in previous experiments and the short duration of our laser ablations, micro-injections or drug treatments might reveal a difference between the establishment and the maintenance of centrosome centration. These two processes are difficult to distinguish experimentally. Furthermore, the specific study of the establishment of centration is challenging because it requires to start from a clearly off-centered position, which is rarely observed in normal conditions. Fortunately, we were able to take advantage of an intermediate stage of the enucleation protocol: just after the centrifugation step, and before the washout of the cytoskeleton drugs that were necessary to weaken cell architecture and allow cell fragmentation, the centrosome was close to cell periphery. By removing all the drugs used during the enucleation process (Cytochalasine D, Y27632, Nocodazole) the centrosome moved away from cell periphery toward the cell center of mass within 4 hours (Figure 4A). Interestingly, by washing out actin drugs only and keeping the Nocodazole, we were able to observe the contribution of the actin network in the absence of microtubules (Figure 4B). Under this condition, we found that centrosome centering was possible but incomplete (13 µm distance to the center as compared to 8 µm in control conditions, Figure 4B). This showed that the actin network is sufficient to provide most of the centration independently of the microtubules.

**FIGURE 4:**
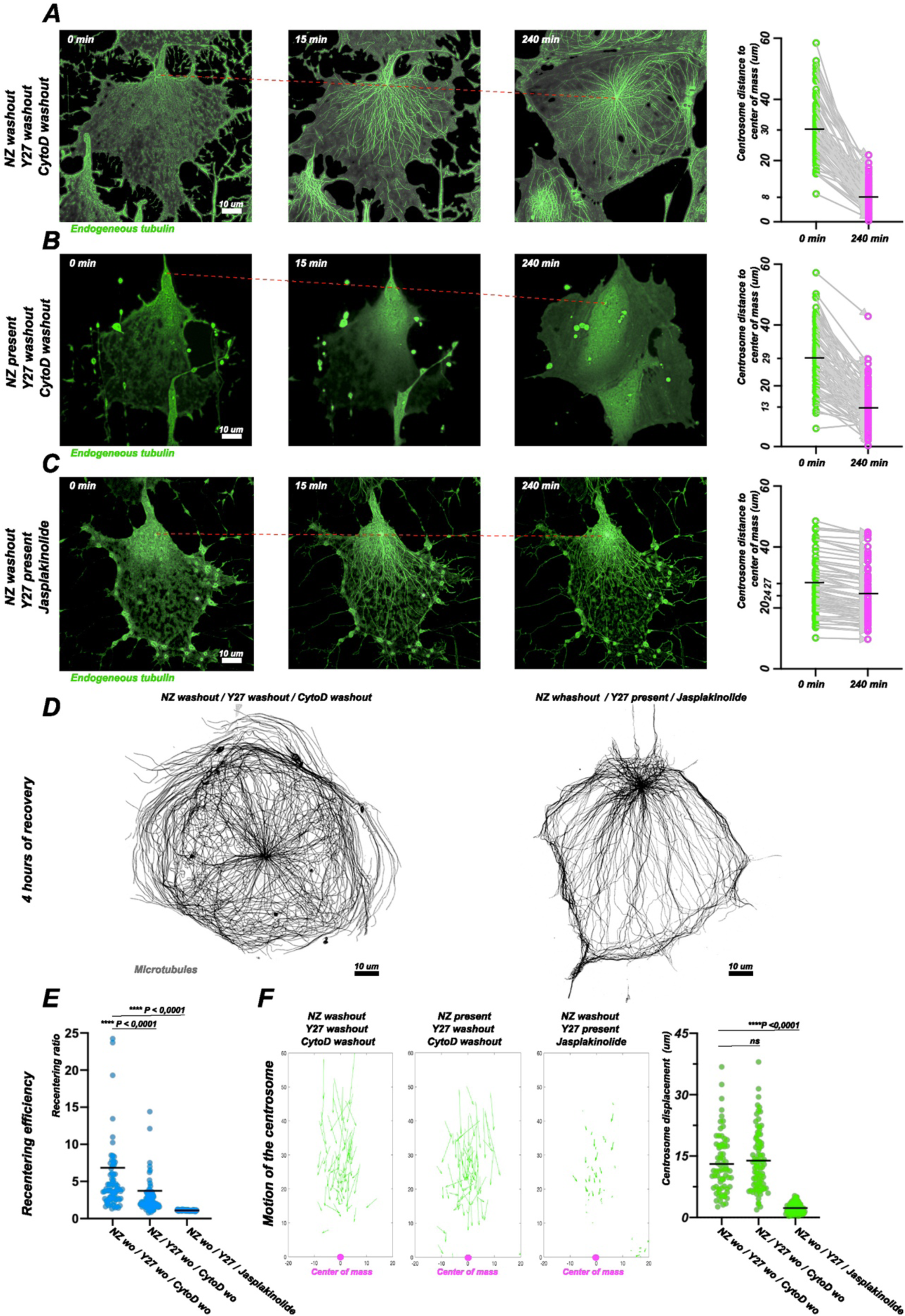
Centrosome recentering following cell enucleation. Briefly, PtK2-GFP cells were enucleated in the presence of Nocodazole (10µM), Y27632 (10µM) and Cytochalasin-D (3ug/ml). After enucleation and before the removal of the enucleation drugs all cytoplasts displayed a pathological actin network with a reproducibly off-centered centrosome biased in the direction of the enucleation pull. To challenge the contributions of actin and microtubules in the recentering processes we designed 3 wash out experiments that were recorded over a period of 4 hours. The control condition consisted in the total wash out of Nocodazole, Y27632 and Cytochalasin-D. To challenge the ability of actin to center a centrosome in the absence of microtubules Nocodazole was kept and only Y27632 and Cytochalasin-D were washed out. To investigate the ability of a dynamic microtubule aster to find the center of a stabilized actin meshwork cytoplasts were first submitted to a 4-hour treatment of Jasplakinolide (600nm) and Y27632 (20µM) in the presence of Nocodazole. Nocodazole was then washed out in the presence of Jasplakinolide and Y27632. In the graphs, horizontal bars represent the mean. P represents the p values, which were obtained from Kruskal-Wallis non-parametric tests. ***(A)*** Image sequence showing microtubule network recovery upon Nocodazole, Y27632 and Cytochalasine-D washout (n=70). The dotted red curve highlights the motion of the centrosome. Histogram shows the distance separating the centrosome and the center of mass before and 4 hours after drugs washout. ***(B)*** Image sequence showing microtubule network recovery upon Y27632 and Cytochalasine-D washout (n=70). Nocodazole is still present. The dotted red curve highlights the motion of the centrosome. Histogram shows the distance separating the centrosome and the center of mass before and 4 hours after drugs washout. ***(C)*** Image sequence showing microtubule network recovery upon Nocodazole washout in the presence of Y27632 (20µM) and Jasplakinolide (600nM) (n=66). Histogram shows the distance separating the centrosome and the center of mass before and 4 hours after drugs washout. ***(D)*** Images illustrate a well-centered microtubule aster 4 hours after Nocodazole, Y27632 and Cytochalasine-D washout (left), and a non-centered aster upon Nocodazole washout in the presence of Y27632 (20µM) and Jasplakinolide (600nM). ***(E)*** Histogram shows the recentering ratio, ie the ratio between final and initial distance separating the centrosome and the center of mass, in the 3 different conditions of centrosome recentering experiments. ***(F)*** Graphs show the displacements of the centrosomes toward the center of mass of the cytoplast in the 3 different conditions of centrosome recentering experiments. Histograms show the total displacement of the centrosomes in the 3 different conditions of centrosome recentering experiments ***(A B C)*** All images were processed following the described pipeline in the dedicated extended material and method section ***(D)*** Images are max projections that were further processed using an unsharp mask and a gamma filter.

Unfortunately, the exact opposite conditions, ie repositioning in the presence of microtubules and in the absence of an actin network, could not be studied. After washing out Nocodazole only, the cytoplasts could not remain spread out and detached rapidly. We therefore designed another approach to assess the ability of a microtubule network to autonomously center a centrosome. By replacing the enucleation buffer with the actin “freezing” buffer immediately after enucleation and washing out the Nocodazole we were able to test the ability of microtubules to recenter the centrosome within a static actin network. Despite normal microtubule regrowth, the aster failed to recenter (Figure 4C). Microtubules span the entire cytoplasm, but the forces produced by their polymerization dynamics, or by molecular motors acting on them, were not sufficient to move the asters against actin filaments, leading to the striking assembly and maintenance of highly off-centered microtubule networks (Figure 4D and S5A).

We then compared the recentering efficiency in these three conditions by measuring the ratio of the distance between the centrosome and the center of mass right after enucleation and 4 hours later. Interestingly, in presence of a dynamic actin network and in the absence of microtubules, ie upon washout of actin drugs but not Nocodazole, recentering occurred but less efficiently than under control conditions (Figure 4E). However, under these conditions of partial recentering, the absolute centrosome displacement was not smaller than in the control condition (Figure 4F). This suggested that microtubules were not required for centrosome displacement, which appeared to be driven by an alternative mechanism based on actin network dynamics. This mechanism was probably non-specific, since it could also center non-functionalized fluorescent beads with the same kinetics and trajectory (Figure S5B).

Careful observation of actin network rearrangements upon washing out of cytoskeleton drugs after cell enucleation revealed that the centrosome centering was not simply driven by centrosome displacement: the cytoplast also moved and reorganized its shape around the centrosome. The centrosome and the center of mass of the cytoplast moved toward one another (Figure 5A). When the cytoplasts changed shape, their center of mass was displaced toward the centrosome (Figure 5B, left). Obviously, when the actin network was “frozen” no rearrangements could occur and the center of mass did not move (Figure 5B, right). However, a dynamic actin network alone was not sufficient to properly drive the reorganization of cell shape. In the absence of microtubules, the displacement of the center of mass was shorter (Figure 5C) and less well oriented than in the control condition with both actin and microtubules (Figure 5D). This prompted us to further quantify the relative contributions of the displacement of the center of mass (ie, the reorganization of the cell shape around the centrosome), and centrosome displacement to the final centering precision (Figure 5E). A first striking observation was that, on average, the motion of the center of mass was responsible for nearly 50% of the centering process (Figure 5F). A closer look at the individual recentering events revealed that smaller distance errors between the centrosome and the center of mass (i.e., good centering events) appeared to be associated with larger displacements of the center of mass rather than the centrosome (Figure 5F). Indeed, the displacement of the centrosome was not correlated to the final error distance (Figure 5G). Unexpectedly, the accuracy of the final centration of the centrosome was rather correlated to the displacement of the center of mass of the cell, ie to the ability of cells to reorganize around the centrosome (Figure 5H). Centrosome centering appeared to be more dependent on the reorganization of the cell around the centrosome than on centrosome displacement itself. In few cases, we could even observe centrosome centering in the absence of any significant centrosome displacement (Figure S5D). Interestingly, experiments in the presence of Nocodazole showed that microtubules were involved in guiding cell reorganization around the centrosome, rather than in the physical movement of the centrosome in the cytoplasm.

**FIGURE 5:**
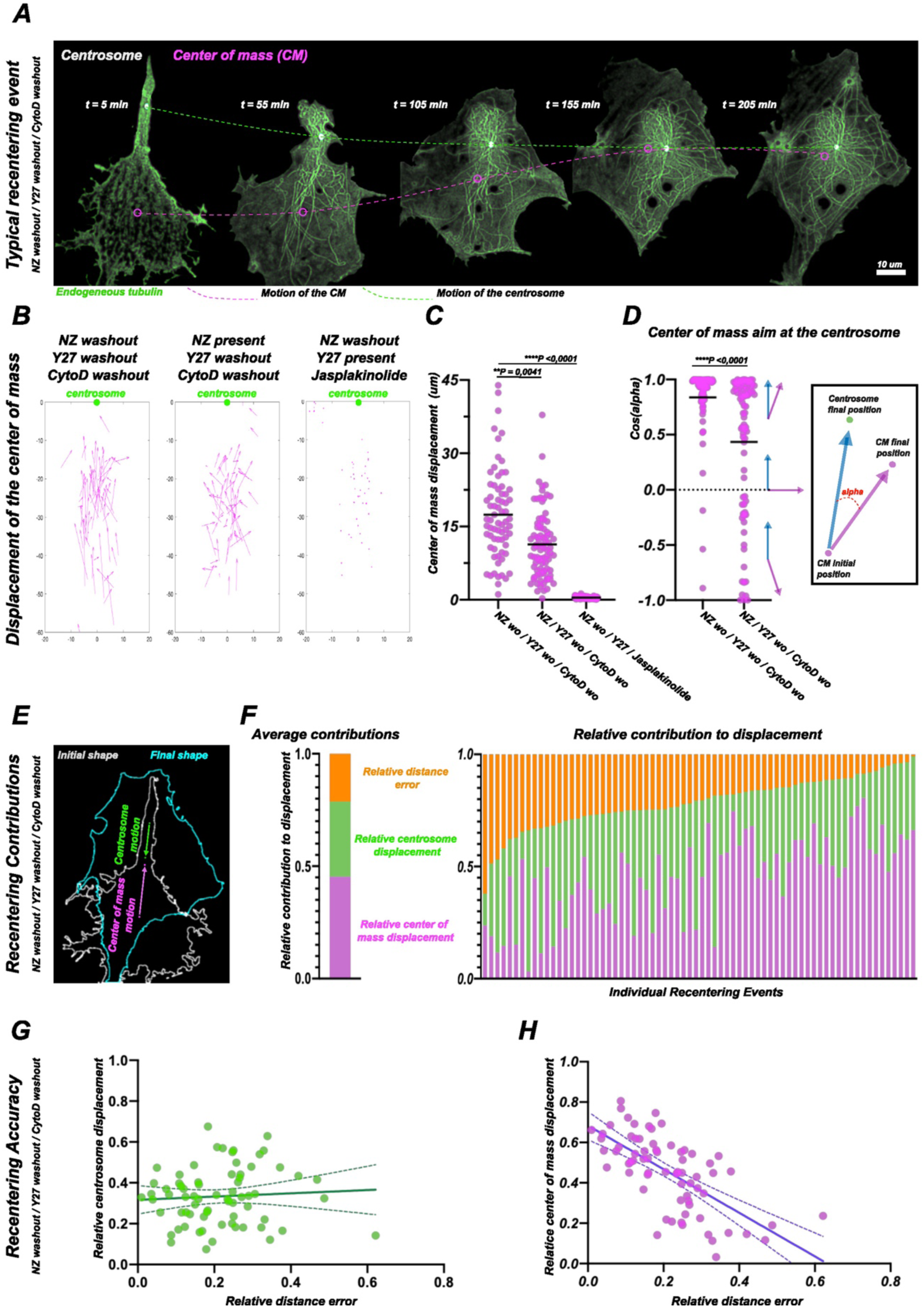
Role of cell shape remodeling in centrosome repositionin. In the graphs, horizontal bars represent the mean. P represents the p values, which were obtained from Kruskal-Wallis non-parametric tests. Nocodazole washout, Y27632 washout, Cytochalasine-D washout (n=70); Nocodazole present, Y27632 washout, Cytochalasine-D washout (n=81); Nocodazole washout, Y27632 present (20µM), Jasplakinolide (600nM) (n=66). ***(A)*** Nocodazole washout, Y27632 washout, Cytochalasine-D washout. Five selected timepoints of a representative live centrosome recentering event showing the repositioning of the centrosome and the center of mass inside a cytoplast over a time course of a few hours. The dotted green line underscores the motion of the centrosome, and the dotted magenta line underscores the motion of the center of mass. ***(B)*** Graphs show the displacements of the cytoplast’s center of mass toward the centrosome in the 3 different conditions of centrosome recentering experiments. ***(C)*** Histograms show the displacement of the center of mass in the 3 different conditions of centrosome recentering experiments. ***(D)*** Histograms show the angle between the axes connecting the initial positions of the centrosome and center of mass and the trajectory of the center of mass. ***(E)*** Overlay of the first and last timepoints of the recentering event depicted in ***(A)***. The arrows represent the magnitude and direction of the displacements of the centrosome and of the center of mass. ***(F)*** Histograms show the relative contributions of the displacements of centrosome and of the center of mass to the recentering event upon washout of all drugs. The « error » corresponds to the final distance between the centrosome and the center of mass. Right histogram shows relative distance measurements in 80 individual recentering events sorted by final error. Left histogram shows their averaged values. ***(G)*** Graph shows the correlation between the centrosome displacement and the distance error. Distances are normalized to the initial distance separating the centrosome and the center of mass. The dark green line represents the best linear fit and the dotted green line represent the fit’s 95% confidence interval. ***(H)*** Same graph as G but showing the correlation between the displacement of the center of mass and the distance error. ***(A)*** Images were processed following the described pipeline in the dedicated detailed material and method section

## Discussion

Taken altogether, these data showed that, in adherent cells in culture, microtubules do not support the propagation of mechanical forces from the cytoplasm to the centrosome. The position of the centrosome appeared to be determined by the architecture and flow of the actin network rather than by a balance of pushing and pulling forces in the microtubule network. This suggested that the organization of the network and the shapes of the microtubules reflect the physical constraints on their growth in the cytoplasm (Brangwynne et al., 2007) rather than the application and long-range transmission of mechanical forces along them. These results render obsolete previous mechanical models of microtubule-based centrosome positioning in adherent cells (Zhu et al., 2010; Letort et al., 2016). On the other hand, they are consistent with the numerous reports showing that microtubules are anchored to actin filaments (Dogterom and Koenderink, 2019) and that the contraction and mechanical properties of the actin network direct and maintain the position of the microtubule aster (Solecki et al., 2009; Chaigne et al., 2016; Colin et al., 2020; Xie et al., 2022). Our findings also indicate that microtubules may serve as sensors and signaling platforms, guiding the reorganization of the cell around the centrosome, rather than purely mechanical players, displacing the centrosome (Verma and Maresca, 2019; Wasteneys, 2004; Parker et al., 2014).

The extent to which microtubule mechanics contribute to the organization of the microtubule network seems to depend on the size and shape of the cell in relation to the organization of the actin network. In adherent cells, microtubules appear embedded inside a dense actin network, which most likely generates a mechanical coupling that prevents autonomous displacements of the microtubule aster. In contrast, in mammalian eggs or mitotic cells, the actin network is more concentrated along the periphery than in the cytoplasm so microtubules are less constrained (Rosa et al., 2015; Tanimoto et al., 2016). These conditions might be more favorable for the propagation of forces along microtubules and the displacement of the aster. According to this view, the degree of freedom of the microtubule aster is inversely correlated to the density of the actin network, and thereby more or less sensitive to external cues. In adherent cells microtubules are constrained by the actin network so the position and asymmetry of the microtubule network aligns with the cellular micro-environment (Jimenez et al., 2021), whereas in mitotic cells, microtubules are more free, so that network architecture is more autonomous and determined by microtubule self-organization (Oriola et al., 2018).

Interestingly, the coupling between microtubules and actin filaments can be modulated and allow the aster to switch between opposite positioning mechanisms and favor aster off-centering. In oocytes, the dense actin network, limits the autonomous centering of the microtubule array and moves the meiotic spindles toward the cell periphery (Schuh and Ellenberg, 2008; Azoury et al., 2008; Chaigne et al., 2013). Conversely, actin disassembly can facilitate centrosome displacement throughout the cytoplasm during ciliogenesis (Kim et al., 2010; Pitaval et al., 2010), as well as during immune synapse (Obino et al., 2015) or axon formation (Solecki et al., 2009). The coupling can also be reduced by several means. The release of microtubules can free the centrosome and allows its translocation by molecular motors (Hannaford et al., 2022). Individual microtubules can also self-organize with molecular motors when they are no longer bound to the centrosome (Rodionov and Borisy, 1997). Indeed, the longitudinal gliding of individual microtubule is less limited by friction than the translocation of an entire aster. Disassembling and reassembling the aster at a different position might be physically easier than displacing it. There are therefore various mechanisms of cytoplasmic clearance or aster disassembly that can circumvent the role of the actin network in controlling microtubule aster positioning.

## Acknowledgements

We thank Jeremie Gaillard, Christophe Guerin and Magali Prioux from the CytoMorpho Lab for the purifications of proteins that were micro-injected in cells. This work was supported by fundings from the Agence Nationale pour la Recherche (ANR grant AAPG2022-PRC-SHARP) and the European Research Council (ERC consolidator grant 771599).

## Author contributions

Alexandre Schaeffer performed all experiments and analyses. Morgan Gazzola and Benoit Vianay provided some guidance on micro-injection and laser nano-ablation. Matthieu Gelin designed the holder for the centrifugation of thin glass coverlsips. Chiara De Pascalis performed preliminary experiments. Laurent Blanchoin and Manuel Théry supervised the project. Alexandre Schaeffer and Manuel Théry designed the study and wrote the manuscript.

## Disclosures

Authors declare that they have no competing interests.

## Materials and Methods

See Supplementary Materials for the references of all reagents.

### Cells culture

PtK2 (Potorous tridactylus kidney) cells (WT and expressing GFP-tubulin) were obtained from Franck Perez lab and were not further characterized. RPE1 (Retineal pigmental epithelial cells) wild-type cells. MEF (Mouse embryonic fibroblasts) cells obtained from the lab of John Eriksson and were not further characterized. Cells were grown at 37°C and 5% CO2 in DMEM/F12 supplemented with 10% fetal bovine serum and 1% AA solution. Upon passage, cells were detached using TrypLE.

### Drug treatments

#### Live actin and microtubule staining

Actin and microtubules were labeled in live using SIR-Actin / SPY FastAct and SPY-Tubulin. The labeling reagents were added directly to the cell culture media 4 hours prior to imaging and were not washed before the acquisitions. SPY-Tubulin and SPY-FastAct were used at a dilution of 1/1000 and SiR-Actin was used at a concentration of 300nM. To further improve the quality of the live stainings we performed all the labeling in the presence of Verapamil (10µM).

#### Actin “freezing”

To fully immobilize the actin network cytoplasts were treated with a combination of Jasplakinolide (600nM) and Y27632 (20µM). The incubation was performed either after the full recovery of the cytoplasts or right after enucleation and lasted for at least 2 hours.

#### Dynein inhibitors

Dynarrestin or Dynapyrazole A were resuspended in DMSO at their solubility limit upon arrival. The drugs were used in the week that followed their resuspension. To inhibit dyneins cells were treated for one hour with either Dynarrestin (50µM) or Dynapyrazole A (80µM) diluted in culture media. Cells were then fixed and stained using a variation of the IF protocol mentioned above (see extended methods for more details).

#### Mini-asters

PtK2-GFP cells were detached and platted on square 20x20 polystyrene slides coated with fibronectin and collagen (both at 20 µg/ml) and placed inside an incubator for at least 24 hours. PtK2-GFP cytoplasts, enucleated in the context of intact microtubule aster, were left 4 hours in the incubator to recover. Cells or cytoplasts were then submitted to various drug treatments for 6 hours prior to fixation and labelling: Nocodazole 6µM withY27632 1µM to obtain high contraction state, Nocodazole 6µM with Y27632 30µM (or 20 µM for cytoplast) to obtain low contraction state, or Nocodazole 6µM with cytochalasin-D 0,5µg/ml to mildly depolymerize actin.

### Cell enucleation

Cells were detached and platted on square 20x20 polystyrene slides coated with fibronectin and collagen (both at 21 µg/ml) and placed inside an incubator for at least 12 hours. Prior to enucleation, cells were incubated 20 min at 37°C in an enucleation buffer (cytochalasin-D, +/- Y27632, +/- Nocodazole). Cells were then enucleated inside the enucleation buffer using high speed centrifugation (13000 RPM) for 25 min at 37°C. Enucleations were performed inside an Avanti JXN-26, Beckman Coulter centrifuge equipped with a swinging rotor (JS-13.1, Beckman Coulter). After enucleation, slides were rinsed with fresh culture media every 3 minutes for 12 minutes. Slides are then placed in the incubator for at least 4 hours to allow a complete recovery of the cytoplasts. Various pretreatment and enucleation buffers were used for the distinct cell types.

For PtK2, cells were submitted to an overnight treatment of Y27632 (5µM). The enucleation buffer contained Cytochalasin-D (3ug/mL) and Y27632 (5µM) (in culture media). To enucleate PtK2 with a mini-aster, cells were submitted to an overnight treatment of Nocodazole (10µM) and Y27632 (10µM). The enucleation buffer contained Cytochalasin-D (3ug/mL), Y27632 (10µM) and Nocodazole (10µM) (in culture media).

To enucleate RPE1 with a mini-aster, cells were not submitted to any overnight treatment prior to enucleation. The enucleation buffer contained Cytochalasin-D (0,65ug/mL), Y27632 (5µM) and Nocodazole (5µM) (in culture media).

To enucleate MEF with a mini-aster, cells were not submitted to any overnight treatment prior to enucleation. The enucleation buffer contained Cytochalasin-D (0,35ug/mL), Y27632 (10µM) and Nocodazole (5µM) (in culture media).

### Centrosome recentering assay

PtK2-GFP cells with endocytosed beads were enucleated in the context of small microtubule asters. Right after enucleation cytoplasts were directly brought to the microscope for imaging. After the first timepoint of the acquisition the enucleation buffer was washed and replaced with fresh culture medium containing additional drugs (Nocodazole (10µM), Jasplakinolide (600nM) and Y27632 (20µM)) in specific cases. To efficiently removed any residual drugs multiple washes were performed. Live acquisitions lasted for 4 hours with a 4µm Z-stack (5 steps spaced by 1µm intervals) every 5 minutes. Two fluorescent channels were recorded: 488nm to follow the endogenous GFP-tubulin and 555nm to follow the motion of the 500nm endocytosed labeled polystyrene beads.

### Immunofluorescence

Cells were fixed 10 min at room temperature in the « cytoskeleton buffer » (MES 10mM, KCl 138mM, MgCl 3mM, EGTA 2mM) supplemented with 10% sucrose, 0.05% Triton-X100, 0.05% glutaraldehyde, and 4% of paraformaldehyde. Aldehyde functions were then reduced using an NaBH_4_ solution (1mg/ml in PBS) for 10 minutes. The samples were then washed 2 times with PBS and 1 time with PBS-Tween 0.1%. The slides were then incubated in a blocking solution (PBS, 0.1% Tween-20, 3% BSA) for 25 min and then with the primary antibodies diluted in the blocking solution for 30 min. The slides were then rinsed 3 times using PBS-Tween 0.1% and incubated in blocking solution for 30 min with the secondary antibodies, and labelled phalloidin. Finally, the slides were rinsed twice in PBS-Tween 0.1%, once in PBS, once in mQ water and were then mounted in Mowiol 4-88.

Microtubules were stained using rat antibodies against tyrosinated tubulin (YL1/2, 1/300). Centrosomes were stained using either mouse antibodies against 𝛾tubulin (1/300) or rabbit antibodies against 𝛾tubulin (1/300) or rabbit antibodies against pericentrin (1/300). The Golgi appartus was stained using either mouse antibodies against GM130 (1/300) or rabbit antibodies against giantin. Actin was stained with Phalloidin labelled with Alexa488 (1/100). The secondary antibodies we used were Donkey anti-rat Alexa 405 (Thermo Fisher, A48268), donkey anti-mouse Alexa 555 (Thermo Fisher, A31570), goat anti mouse Alexa 405 (Thermo Fisher, A31553), donkey anti rabbit Alexa 647(Jackson Immuno Research, 711-605-152). The 74.1-antibody which was micro-injected in cells was detected using a secondary anti mouse antibody. When necessary, sequential stainings with primary and secondary antibodies were performed in order to use several mouse primary antibodies on the same cells.

### Slides coating and patterning

#### Cleaning

Glass slides were submitted to 3 successive washing steps. First, slides were sonicated in acetone for 20-30 min. Then slides were sonicated in pure isopropanol for 20-30 min. To finish slides were sonicated in milliQ water for 20-30 min before being air dried.

#### Polystyrene coating

To promote cellular adhesion glass slides were coated with a very thin layer of polystyrene. Clean slides were activated using a 3 min air plasma treatment (PE-50 plasmatech). The slides were then placed inside a semi-closed recipient heated at 75°C in the presence of a few drops of hexamethyldisilazane (HMDS) for at least 6 hours. After HMDS treatment the slides were recovered, and spin coated with polystyrene. Briefly, the slides were mounted inside a spin coater (WS-650m2-23NPPB, Laurell) their surface was covered with a solution of 1% polystyrene (the polystyrene is dissolved in toluene) and were subsequently spined for 30 seconds at 1500 rpm. After spin coating the slides were collected dried and stored for a least 1 hour prior to their use.

#### Protein coating

Polystyrene-coated slides were activated using air plasma treatment (PE - 50 plasmatech) for 45 seconds. After activation the slides were then placed inside a solution of proteins (12 or 21 µg/ml of fibronectin and collagen diluted in sterile PBS) for at least 30 minutes at room temperature. Slides were rinsed twice in milliQ water and once in sterile PBS.

#### Surface passivation

Polystyrene-coated slides were activated using air plasma treatment (PE - 50 plasmatech) for 45 seconds and immersed in a solution of 0.1 mg/ml of poly(L-Lysine)- poly(ethylene-glycol) (PLL-PEG JenKemTechnology) in 10 mM Hepes (pH 7.4) for 1 hour. PEGylated slides were briefly plunged into milliQ water and were rapidly dewetted. The PLL-PEG slides were then stored at 4°C for at least 1 hour.

#### Micropatterning

PEGylated coverslips were put in tight contact with a quartz-chrome printed photomask (Toppan Photomask). Tight contact was maintained using a vacuum holder. The PEG layer was burned with deep UV (190nm) through the non-chromed windows of the photomask, using a UVO cleaner (Model No. 342A-220, Jelight), at a distance of 1cm from the UV lamp with a power of 6mW/cm2, for 5 min. After exposure, the patterned PLL-PEG coated slides were gently detached from the mask by submerging them under milliQ water. Once the slides detached, they were dewetted and incubated in a solution of adhesion protein according to the protocol for protein coating.

### Micro-injection

Cells were detached and platted on polystyrene slides coated with fibronectin and collagen (both at 21 µg/mL) and placed inside an incubator for at least 12 hours. Alternatively, cells were detached, and platted on polystyrene patterned slides coated with fibronectin and collagen (both at 12 µg/mL) and placed inside an incubator for 12 hours. Right before injection, culture medium was exchanged to remove dead cells and floating debris. Glass microneedles were pulled from clark borosilicate thin wall capillary (30-0050Harvard Apparatus) using a vertical pipet puller (PC-100, Narishige). Microneedles were manually controlled with an InjectMan 4 micromanipulator (Eppendorf). Microinjection of cells were performed on an inverted microscope (Nikon Ti2 Eclipse) equipped with a Prime BSI Express CMOS camera (Photometrics) and using a Nikon CFI Plan Fluor 40x/0.75 NA dry objective. Various compensation pressures were applied using a FemtoJet 4i (Eppendorf) pump. Cell medium was maintained at 37°C during the whole experiment using a H-301 heating chamber (Okolab). Micro-Manager 1.4.21 software was used for live images acquisition during the micro-injection procedure. After micro-injection, cellular media was exchanged, and cells were placed inside an incubator for a recovery period.

### Tubulin speckles

Purified labelled tubulin was thawed on ice and diluted in injection buffer (50 mM potassium glutamate, 1 mM MgCl2, pH 6.8) to obtain a final concentration of 1µM. The obtained micro-injection tubulin solution was kept on ice and protected from light. Microneedles were loaded with 10µL of micro-injection tubulin solution and connected to the Femtojet system. The compensation pressure used was set between 30 to 35 hPa. After micro-injection, cellular media was exchanged, and cells were placed inside an incubator for a 2 hour recovery period.

### Actin speckles

Purified labelled actin was diluted in injection buffer (50 mM potassium glutamate, 1 mM MgCl2, pH 6.8) to obtain a final concentration of 1µM. The obtained micro-injection actin solution was kept on ice and protected from light. Microneedles were loaded with 10 µL of micro-injection solution and were connected to the Femtojet system. The compensation pressure used was set between 70 to 90 hPa. After micro-injection, cellular media was exchanged, and cells were placed inside an incubator for a 2 hour recovery period.

### Antibody against dyneins

Microneedles were loaded with 2µl of the 74,1-antibody commercial solution and were connected to the Femtojet system. The compensation pressure used was set between 25 to 35 hPa. Cells were injected for 15 min and then placed inside the incubator for an incubation and recovery period of 50 min. After the incubation period cells were fixed according to the immunofluorescence protocol. Injected cells were identified using a secondary anti-mouse antibody that recognized the 74,1 antibody.

### Microscopy

Acquisitions were made with a confocal spinning disk microscope (Nikon Ti Eclipse equipped with a spinning scanning unit CSU-X1 Yokogawa) and a R3 retiga camera (QImaging). Images were acquired using either Nikon CFI Plan Fluor 40x/1.3 NA oil objective, Nikon Plan Apo VC 60x/1.40 NA oil objective, Nikon CFI Super Fluor 100x/1.3 NA oil objective. Each wavelength was acquired separately. Metamorph software was used for images acquisition. Live acquisitions were performed inside a live module (kept at 37°C and 5% CO2) mounted on the confocal spinning disk microscope. In nearly all live acquisitions, HEPES (1M PH 7.4) (1/100) and Oxyfluor (1/100) were added to the culture media.

### Laser induced photodamage

Photoablation was performed on the aforementioned spinning-disc system from Nikon using the iLas2 device (Gataca Systems) equipped with a passively Q-switched laser (STV-E,ReamPhotonics) at 355 nm producing 500 picosecond pulses. Laser displacement, exposure time and repetition rate were controlled via ILas software interfaced with MetaMorph (Universal Imaging Corporation). Laser photoablation and subsequent imaging were performed with a CFI Super Fluor 100X/1.3 NA oil objective. Damages to the microtubule and actin networks were performed in parallel to image acquisition using the single spot ON-FLY function. This allowed a high flexibility and live definition of the localizations of laser ablation events. The laser power used was set to 15% and the spot length was set to 500. Consequences of microtubule network ablations on centrosome position were monitored by recording a 2µm Z-stack around the focal plane of the centrosome every 10 seconds.

### Image processing and data analysis

Most image processing and analysis were performed using the Fiji software. Additional processing and analysis were performed on MATLAB. Graph design and statistical analysis were done on GraphPad PRISM. See Supplementary Methods for detailed image analysis protocols.

## Notes

### Competing Interest Statement

The authors have declared no competing interest.

