## supplementary information for "Centrosome positioning independently of microtubule-based forces"

#### Supplementary Materials (pages 2-3)

#### Supplementary Methods (pages 4-6)

#### Supplementary Figures S1 to S5 (pages 7-16)

#### Supplementary Movies S1 to S9 (page 17)

### Supplementary Materials

**Cells and cell culture**

PBS (Gibco, 14190169)
DMEM F12 (Gibco, 31331028)
FBS (Life Technologies, 10270106)
AA solution (Gibco,15240062).
TrypLE (Gibco, 12605036)
Red carboxylate fluorospheres (Thermo fisher, F-8812)

**Slides coating**

Polystyrene (Acros Organic, MW 260,000, 178891000)
Toluene (Sigma, 179418)
HMDS (sigma, 440191)
Fibronectine (Sigma, F1141)
Rat tail collagen I (Gibco, A1048301)
Pll-PEG (JenKem, PLL20K-G35-PEG2K)

**Immunofluorescence**

MES (sigma, M3671)
MgCL2 (Sigma, 208337)
KCl (Sigma, P3911)
EGTA (Sigma, E3889)
PFA (EMS, 15710),
Glutaraldehyde (Sigma, G5882),
NaBH4 (sigma, 71320)
Triton X100 (Sigma, T8787)
Tween (Sigma,1379)
BSA (Sigma, A2153)
Mowiol (Sigma, 81381).
Rat YL1/2 (Merck, MAB1864)
Mouse anti-gamma tubulin (Sigma, T6557)
Rabbit anti-gamma tubulin (Invitrogen, MA52514)
Rabbit anti-pericentrine (Abcam, ab4448)
Mouse anti-GM130 (Fisher Scientific, 610823)
Rabbit anti-giantin (Abcam, ab80864)
Donkey anti rat Alexa 405 (Thermo Fisher, A48268)
Donkey anti mouse Alexa 555 (Thermo Fisher, A31570)
Goat anti mouse Alexa 405 (Thermo Fisher, A31553)
Donkey anti rabbit Alexa 647(Jackson Immuno Research, 711-605-152)
Phalloidin Alexa488 (Life Technologies, A12380)

**Live actin and microtubule staining**

SIR ACTIN (Tebubio, SC001)
SPY-FASTACT-555 (Tebubio, SC205)
SPY-FASTACT-650 (Tebubio, SC505)
SPY-TUBULIN-555 (Tebubio, SC203)
Verapamil (Tebubio, SC001)
HEPES (Sigma, H3375)
Oxyfluor (Sigma, SAE0059)

**Drugs and inhibitors**

Cytochalasin-D (Sigma, C8273)
Jasplakinolide (Sigma, J4580)
Y27632632 (Sigma, Y0503)
Nocodazole (Sigma, M1404)
Dynarrestin (Sigma, SML2332)
Dynapyrazole A (Sigma, SML2117)
74,1-antibody (Sigma, MAB1618)
DMSO (Sigma, 276855)

### Supplementary Methods

#### Proteins purification and labelling

Tubulin was purified from fresh bovine brain using three cycles of temperature-dependent assembly/disassembly in Brinkley Buffer 80 (BRB80: 80 mM Pipes pH 6.8, 1 mM EGTA and 1 mM MgCl2)(Vantard et al., 1994). MAP-free neurotubulin is subsequently purified by cation exchange chromatography (EMD SO, 650 M, Merck) in 50 mM Pipes, pH 6.8, supplemented with 0.2 mM MgC12, and 1 mM EGTA. Fluorescently labelled tubulin (ATTO-565) was prepared by the following previously published method (Shelanski, 1973). Labeled tubulin later used for cellular micro-injection is stored at -80°C inside micro injection buffer (50 mM potassium glutamate, 1 mM MgCl2, pH 6.8).

Actin was purified from rabbit skeletal-muscle acetone powder (Spudich and Watt, 1971). Monomeric Ca-ATP-actin was purified by gel-filtration chromatography on Sephacryl S-300 at 4°C in G buffer (2 mM Tris–HCl, pH 8.0, 0.2 mM ATP, 0.1 mM CaCl2, 1 mM NaN3 and 0.5 mM dithiothreitol (DTT)). Actin was labelled on lysines with Alexa-488 succimidyl ester (Molecular Probes). Monomeric labeled actin later used for microinjection was typically stored concentrated (around 20 - 40µM) and highly labelled (around 70%) inside 60% glycerol G-Buffer at -20°C.

#### Beads endocytosis

20 µl of 500nm red fluorospheres solution was pipetted and centrifuged for 15 min at 13000g at room temperature inside an Eppendorf microcentrifuge equipped with an FA-24 rotor. After centrifugation, the supernatant was discarded, and the bead pellet was resuspended using 1 ml of culture medium. Classically, PtK2 cells were cultivated in T-75 culture flasks filled with 10 ml of culture media. When cells were cultivated in the presence of beads, 500 µl of the resuspended bead solution was added to 9.5 ml of culture media. Cells were kept for a day or two in the presence of the beads to ensure sufficient endocytosis before the experiments.

#### Image processing and analysis

All temporal sequences were aligned with the Fiji plugin “StackReg” prior to further processing.

##### Centrosome detection and tracking

In fixed cells, Z-stacks of $\gamma$tubulin immunostaining were projected in a single plane using the « maximum projection » of Fiji and centrosome position was further extracted by using the « find maxima » plugin. In living cells, temporal acquisitionsof Z-stacks of GFP-tubulin or SIR-tubulin were projected using the « maximum projection » of Fiji and further processed using a gaussian filter (sigma = 2pixels). The centrosomal area was first manually delimitated and thresholded. The centroid of this area was defined as the position of the centrosome.

Centrosome tracking was performed using the « trackmate » plugin of Fiji. Tracks were constructed using the simple LAP tracking algorithm. Tracking parameters were finely tuned for each centrosome tracking event. In ablation experiments, centrosome displacements away from the ablated region were counted positively. Temporal projection of centrosomal absolute displacements on a single axis are performed using a MATLAB homemade macro.

For the vectorial displacement map of the centrosomes, all the images were realigned on the final position of the center of mass. The graphs were plotted using the MATLAB « quiver plot » tool and the vector were scaled by a factor of 0,5.

**Microtubule network analysis**

Microtubule amount was assumed to scale with the total fluorescence intensity of GFP-tubulin along the microtubule network. The microtubule network was first segmented on the first timepoint of the acquisition to create a mask of the microtubule network. This mask was subsequently projected onto a single plane. The intensity of the positive pixels was set to be equal to one and the negative pixels are set to be equal to zero. Z-stacks of the tubulin signal were projected in a single plane using the « SUM projection » of Fiji. The mask was then multiplied with these images to measure the total pixel intensity.

In ablation experiments, only the microtubules on the side of the centrosome where the ablation occurred were considered. See the graphical representation in Figure S1 for the geometrical definition of the microtubule network disruption region and its quantification. The total intensity along a line crossing centrosomal microtubules represents the initial amount of microtubule on this side of the centrosome prior to laser ablation. Along this same line the total intensity of ablated microtubules was computed. The ratio of the total fluorescence of ablated microtubules over the total fluorescence of all considered microtubules yielded the microtubule depletion ratio.

**Golgi apparatus analysis**

Z-stacks of the GM130 immuno-staining were projected in a single plane using the « MAX projection » of Fiji. The position and the structure of the Golgi were first manually thresholded to generate binary masks for further automated analysis. Manualy thresholded masks of the Golgi apparatus were processed using the « remove outlier » and « maximum filter » of Fiji. The modified images generated from the Golgi masks were then automatically thresholded using the max entropy method. A final step of binary processing was used to fill the holes inside the newly created masks. This allowed the generation of almost continuous Golgi apparatuses, the outlines of which were defined using the « create selection » function of Fiji. The area enclosed inside this outline curve was further measured using the « measure » function of Fiji and was interpreted as the size of the Golgi apparatus (see Figure S3).

To quantify the fragmentation of the Golgi apparatus, the masks were used to extract the amount and size of fragments using a combination of the « analyze particle » and the « measure » functions of Fiji.

**Actin and tubulin speckles detection and tracking**

First, speckle images were segmented using the "subtract background" function of Fiji. Then the « normalized local contrast » integral filter was applied. To remove the additional noise created by preceeding filter, we applied a « gaussian blur filter » coupled to a manual intensity subtraction. Finally, another gaussian blur filter was applied to quench the remaining noise (Figure S2).

Speckles in segmented images were tracked using the « trackmate » plugin of Fiji. Tracks were constructed using the « simple LAP tracking » algorithm (Figure S2). Tracking parameters were finely tuned for each tracking event. A total period of 15 or 30 seconds centered on the ablation event was defined to analyze the behavior of the actin filaments or microtubules. The periods before and after ablation were each composed of 6 frames and shared a common frame (the frame right before ablation). Only tracks lasting for the entire period were considered. The total displacements of the tracks were extracted and were averaged.

The reconstituted speckle tracks were imported into MATLAB to compute the displacement vectors between the first and last positions of the speckle in each individual tracks. The displacement vectors were then plotted using the « quiver plot » tool.

**Cell shape analysis**

Cells or cytoplasts boundaries were defined using the staining of the actin network, or the GFP-tubulin signal in live cell experiments. Z-stacks fluorescent signal were projected in a single plane using the « max projection » of Fiji. The contour of the cell was extracted through manual thresholding of the image. The area and position of the centroid, ie center of mass of the signal in the segmented region, were computed using the « measure » function of Fiji.

For the vectorial displacement map of the centroids, all the images were realigned on the final position of the centrosome. The graphs were plotted using the MATLAB « quiver plot » tool and the vectors were scaled by a factor of 0,4.

### Supplementary Figures

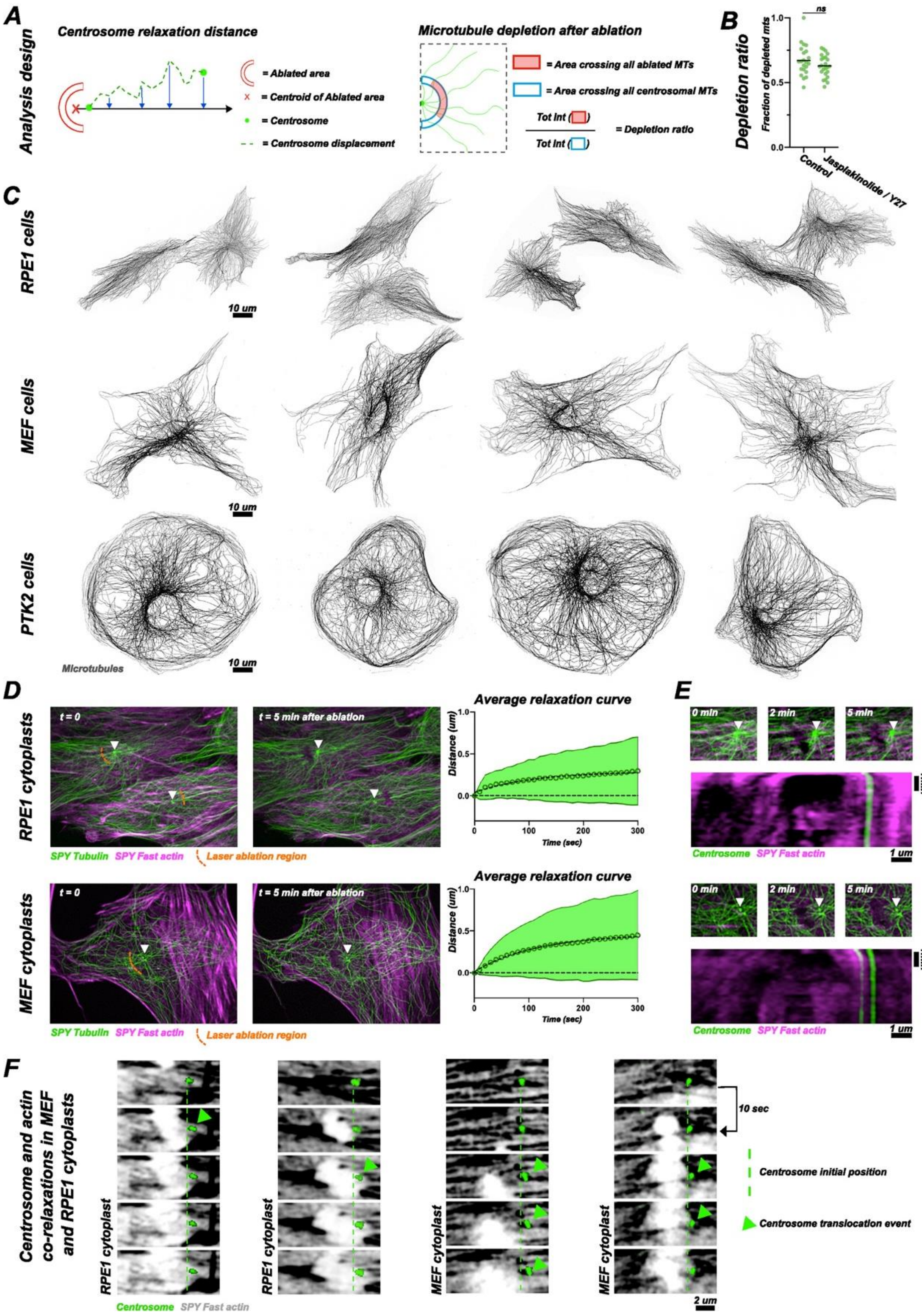


**FIGURE S1: Microtubule network relaxation upon ablation in RPE1 and MEF cells**

***(A)*** Schematic representation of the quantification of centrosome relaxation (left) and of the fraction of microtubule depletion ratio (right) upon microtubules ablation.

***(B)*** Histogram shows the the proportion of depleted microtubules on the side of the centrosome where the laser ablations were performed in the control condition (n=22) and in the case of Jasplakinolide (600nM) and Y27632 (20µM) treatment (n=24). Horizontal bars represent the mean. P values were obtained from Kruskal-Wallis non-parametric tests.
***(C)*** Images of the microtubule network of RPE1 cells, MEF cells and PtK2 cells.
***(D)*** Relaxation of the actin network (magenta) and microtubule network (green) upon repeated laser ablations during five minutes in RPE1 cytoplasts (top) and in MEF cytoplasts (bottom). Images show images of the first and last timepoints of a laser ablation experiment. White arrowheads indicate the positions of the centrosomes. Graphs show the mean relaxation profile of the centrosomes (30 RPE1 cytoplasts and 36 MEF cytoplasts). Circles represent the average displacement of the centrosome at each timepoint, and the continuous lines represent the standard deviation.
***(E)*** Magnified view of the centrosomal area of a RPE1 cytoplast (top) and a MEF cytoplast (bottom) depicted in D. White arrowheads indicate the position of the centrosome. Kymographs show the centrosome and actin relaxations in the zoomed in area. The linescans (scaled 3 times to smoothen the signal) were performed along a line connecting the centrosomes with the centroids of the ablated areas and spanning the entire length of the zoomed in areas.
***(F)*** Magnified views showing actin network (grey) and the centrosome (green) during the repeated laser ablation experiment in RPE1 and MEF cytoplasts. The temporal sequences are made of consecutive images covering a total period of 40 seconds. Small centrosomal translocation events can be visualized in parallel to local relaxations of the surrounding actin meshwork.

***(C D)*** All images are max projections, further processed using an unsharp mask, and a gamma filter.
***(F)*** All images are max projections, further processed using a gaussian blur filter. The LUT of the SPY FAST actin were inverted to facilitate the visualization of the local destruction and relaxation of the meshwork.

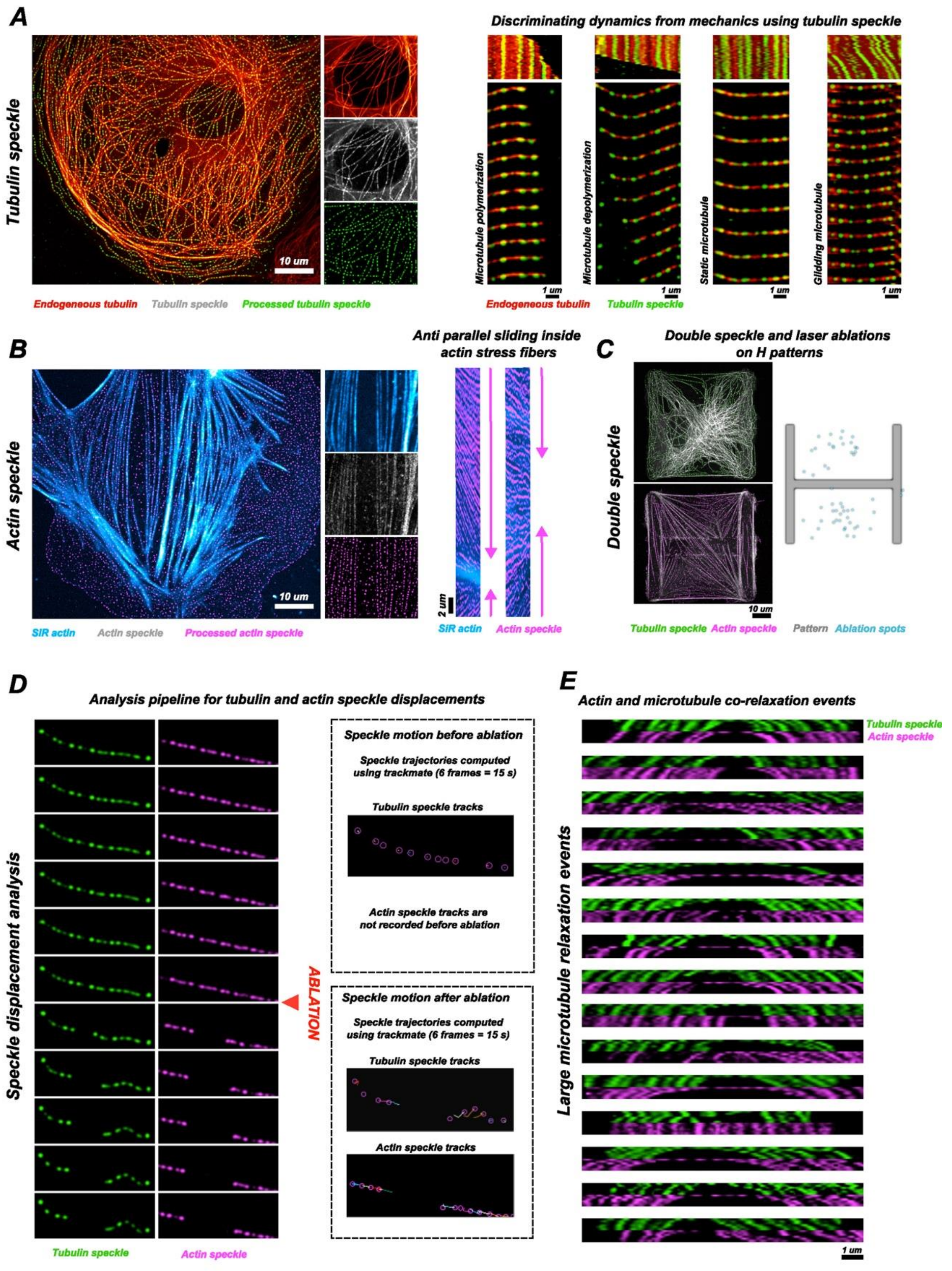


**FIGURE S2: Double speckle microscopy and speckle tracking analysis**

***(A)*** PtK2 cell expressing tubulin-GFP (red), 2 hours after micro-injection of 1µM of purified Atto-555-tubulin (green). On the left, images show the microtubule network with the non-processed (grey) and processed (green) tubulin speckles along the microtubules. On the right, kymographs (constructed over a period of 108s) along the length of a few selected microtubules. Selected timepoints extracted from the video used to build the kymograph can be found below each corresponding kymograph. Selected sequences show a polymerizing microtubule, a depolymerizing microtubule, a static microtubule, and a gliding microtubule . Kymographs are scaled by a factor of 3 for smoothness.

***(B)*** WT PtK2 cell live-stained using SIR-Actin, 2 hours after micro-injection of 1µM of purified Alexa-488-actin. Image shows the actin network with the non-processed (grey) and processed (blue) actin speckles along the actin structures. Kymographs (constructed over a period of 30 minutes) along the length of two selected stress fibers. Selected sequences show contraction events with the presence of antiparallel speckle slidings along the length of both stress fibers. Kymographs are scaled by a factor of 3 for smoothness.

***(C)*** Position of laser ablations in PtK2 cells platted on 3500um$^2$ patterns and co-injected 2 hours earlier with purified and labelled actin and tubulin (1µM and 1.5µM respectively). Images show the tubulin (top) and actin speckles (bottom). The map shows the localizations of all the analyzed ablation events.

***(D)*** Analysis pipeline used to describe the behavior of a microtubule undergoing ablation. Briefly, the motion of the tubulin speckles were tracked 15 sec before ablation and then again for 15 sec after ablation using the simple LAP tracking method in Trackmate. When microtubules displayed significant relaxations, the motions of the surrounding actin speckles were tracked in parallel to that of the tubulin speckles. The amplitudes of the displacements of individual speckles were averaged to yield the mean amplitude of the displacement of the microtubule before and after ablation and the mean amplitude of the displacement of the surrounding actin meshwork after ablation.

***(E)*** Large microtubule relaxation events (motions > 400 nm, depicted in fig 1 G). Kymographs show the coordinated motions of the actin and tubulin speckles in the 15 seconds following laser ablation. The kymographs are constructed along the length of the moving microtubules and are scaled by a factor of 3 for smoothness.

***(A B C D E)*** *For* the details regarding actin and tubulin speckle processing, see the dedicated section in the extended material and methods.

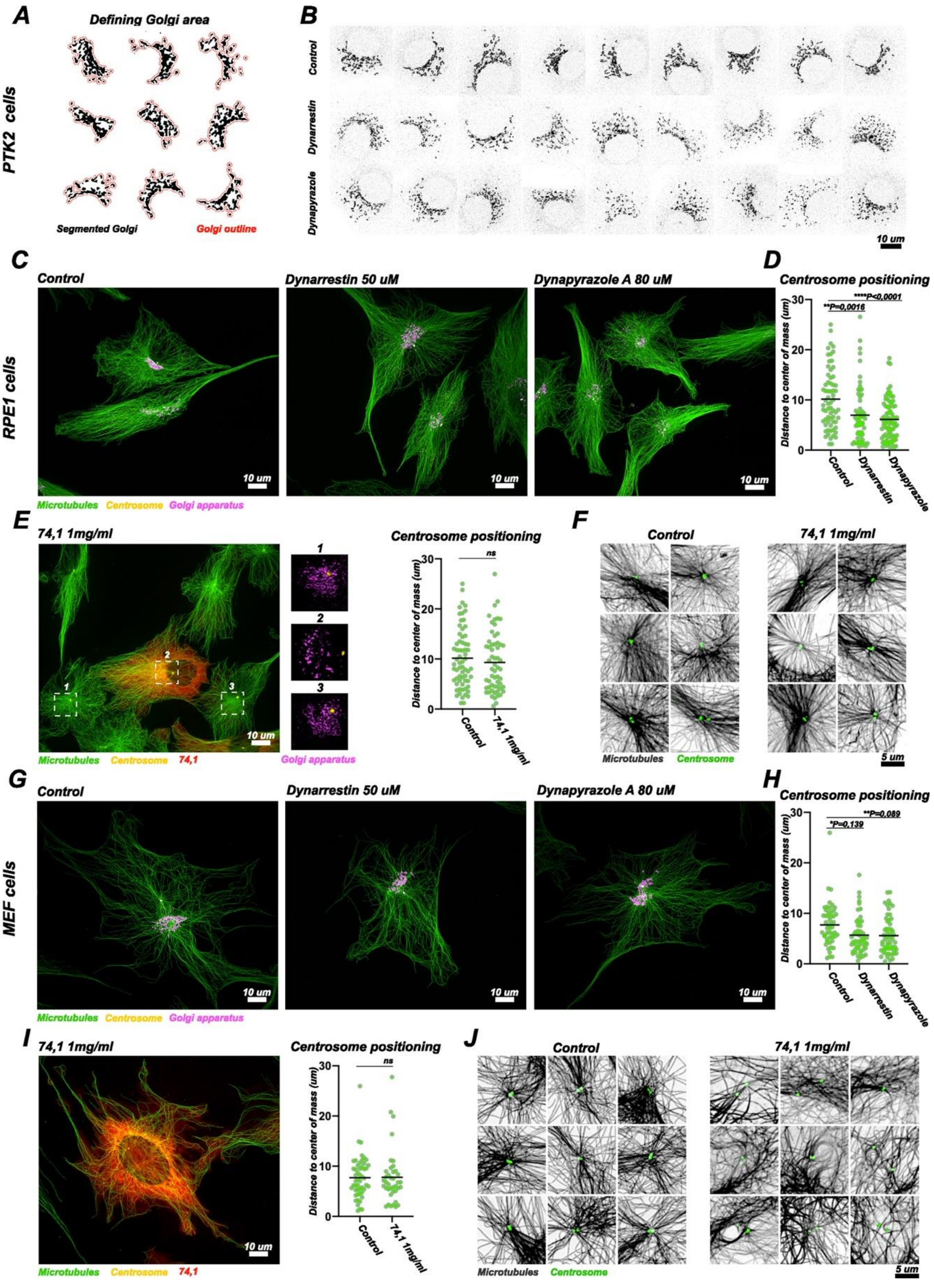


**FIGURE S3: assessing the effect of dynein inhibition in MEF and RPE1 cells**

The activity of dynein molecular motors was inhibited inside RPE1 and MEF cells using a one-hour treatment of Dynarrestin (50µM), Dynapyrazole A (80µM) or through the micro-injection of the 74.1 (1mg/ml) inhibiting antibody. After micro-injection cells underwent a 50-minute incubation period. In all conditions cells were fixed and stained for the microtubules, the centrosome, the Golgi apparatus, and actin. In the graphs, horizontal bars represent the mean. P represents the p values, which were obtained from Kruskal-Wallis non-parametric tests.

***(A)*** Illustration showing nine segmented Golgi apparatuses from the PtK2 control condition and their outlines. The size of the Golgi apparatus is computed as the area enclosed by the curve defined as the Golgi outline (see extended material and methods for the processing used to define the outline curve of the Golgi apparatus).
***(B)*** Images show examples of Golgi apparatuses in the control PtK2 cells (top), upon one hour of Dynarrestin treatment (middle) and one hour of Dynapyrazole A treatment (bottom).
***(C)*** Images show examples of Golgi apparatuses and microtubules in the control RPE1 cells (left), after one hour of dynein inhibition using Dynarrestin (middle), or Dynapyrazole A (right).
***(D)*** Histogram shows the distance between the centrosome (whose position was determined using the gamma tubulin staining) and the center of mass of the cell (whose periphery was determined using the actin staining) in RPE1 cells, in the control (n=66), Dynarrestin (n=60) and Dynapyrazole A (n=83) conditions.
***(E)*** The image shows a RPE1 cell fixed 50 minutes after micro-injection with the 74.1 (1mg/ml) antibody. Graph shows the distance between the centrosome (whose position was determined using the gamma tubulin staining) and the center of mass of the cell (whose periphery was determined using the actin staining) in the control (n=66) and 74,1 (n=59) micro injections conditions.
***(F)*** Images show examples of pericentrosomal microtubule network in RPE1 cells in control conditions (left) and 50 minutes after the micro-injection with the 74.1 antibody (right).
***(G)*** Images show examples of Golgi apparatuses and microtubules in the control MEF cells (left), after one hour of dynein inhibition using Dynarrestin (middle), or Dynapyrazole A (right).
***(H)*** Histogram shows the distance between the centrosome (whose position was determined using the gamma tubulin staining) and the center of mass of the cell (whose periphery was determined using the actin staining) in MEF cells, in the control (n=50), Dynarrestin (n=57) and Dynapyrazole A (n=52) conditions.
***(I)*** The image shows a MEF cell fixed 50 minutes after micro-injection with the 74.1 (1mg/ml) antibody. Graph shows the distance between the centrosome (whose position was determined using the gamma tubulin staining) and the center of mass of the cell (whose periphery was determined using the actin staining) in the control (n=50) and 74,1 (n=39) micro injections conditions.
***(J)*** Images show examples of pericentrosomal microtubule network in MEF cells in control conditions (left) and 50 minutes after the micro-injection with the 74.1 antibody (right).

***(B C E F G I J)*** All images are max projections, further processed using an unsharp mask, and a gamma filter.

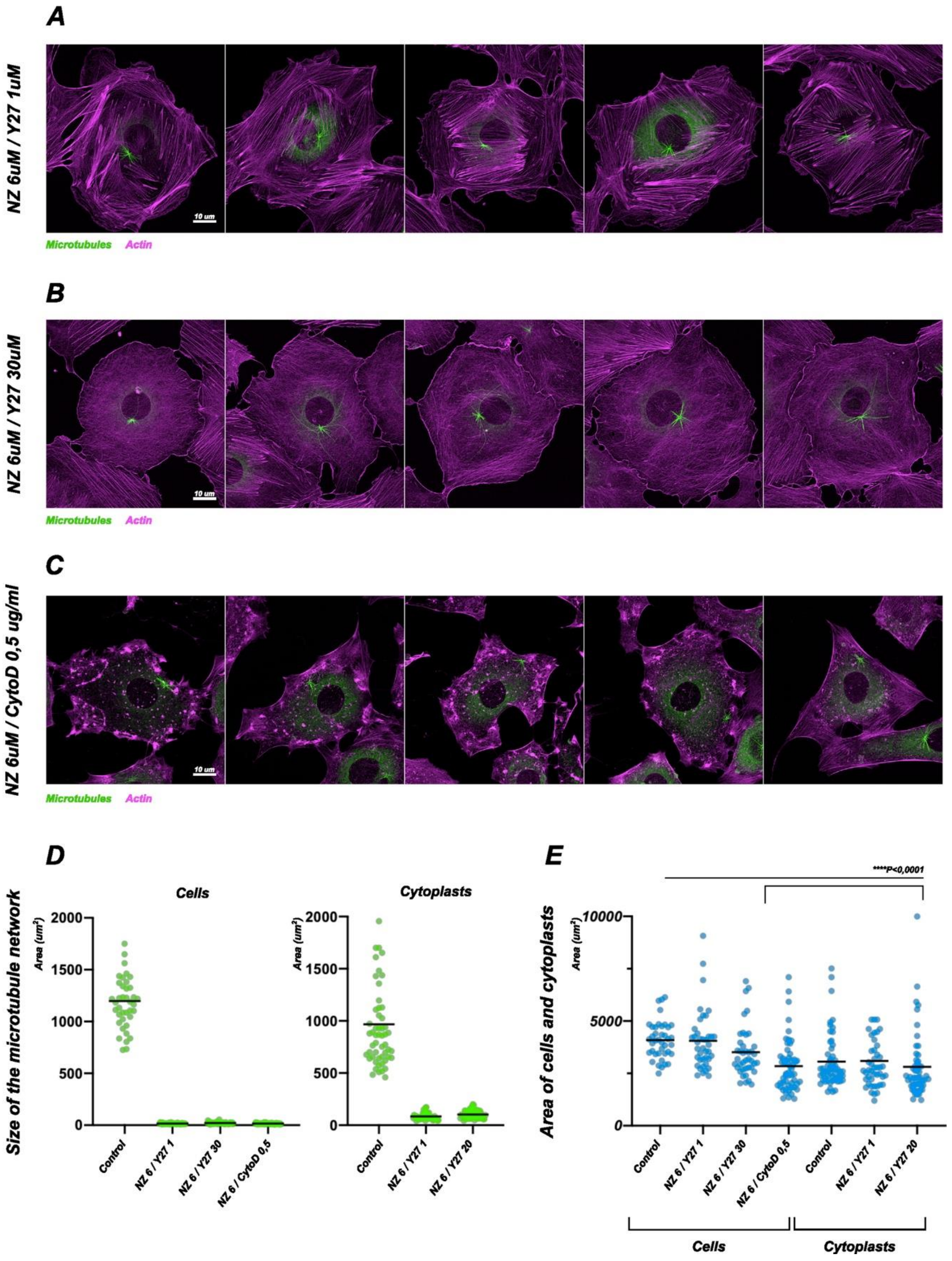


**Figure S4: Regulation of the position of mini-asters of microtubules.**

Mini-asters were induced in PtK2 cells and cytoplasts. Briefly, cells or cytoplasts were exposed to 6 hours of Nocodazole and Y27632 or Cytochalasine-D at various doses to generate small microtubule asters inside strongly contractile, poorly contractile, or disrupted actin networks. In the graphs, horizontal bars represent the mean. P represents the p values, which were obtained from Kruskal-Wallis non-parametric tests. For each condition, at least 40 different cells or cytoplasts were analyzed.

***(A)*** Representative fixed images of well centered mini-asters in highly contractile cells (Nocodazole (6µM), Y27632 (1µM)).
***(B)*** Representative fixed images of well centered mini-asters in poorly contractile cells (Nocodazole (6µM), Y27632 (30µM)).
***(C)*** Representative fixed images of off-centered mini-asters upon actin network disruption (Nocodazole (6µM), Cytochalasin-D (0,5 ug/ml)).
***(D)*** Graphs representing the size of the microtubule network in the various cells (left) and cytoplasts (right). The size of the microtubule network was defined as the projected area occupied by the microtubule network (see the dedicated section inside the extended material and methods).
***(D)*** Graphs representing the spreading area of the various cells (left) and cytoplasts (right). The spreading area was determined using the actin channel (see the dedicated section inside the extended material and methods)

***(A B C)*** All images are max projections, further processed using an unsharp mask and a gamma filter.

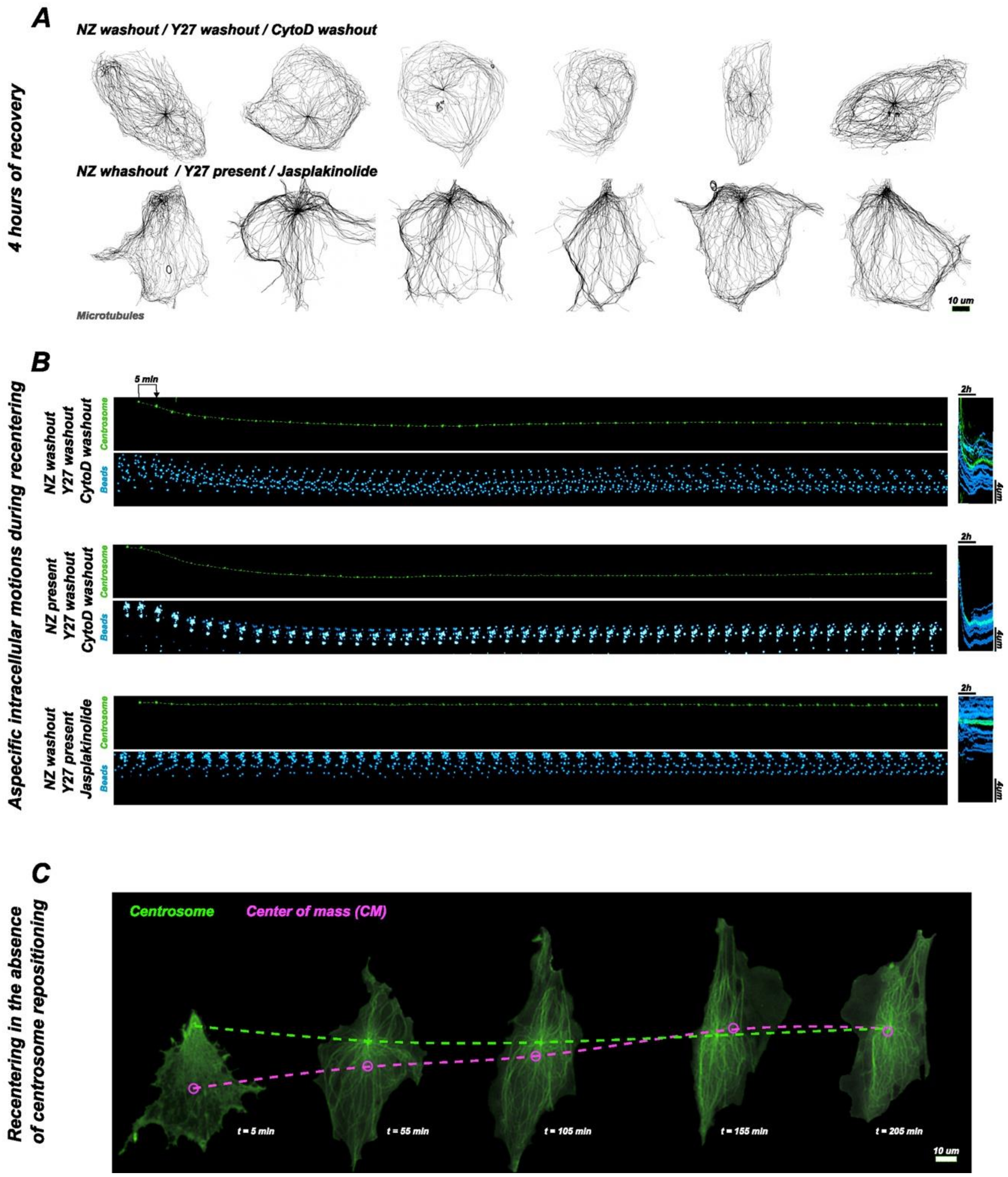


**FIGURE S5: centrosome recentering following cytoplast enucleation**

PtK2 cells stably expressing tubulin-GFP were enucleated in the presence of Nocodazole (10µM), Y27632 (20µM) and Cytochalasin-D (3ug/ml). To challenge the contributions of actin and microtubules in the recentering process we designed 3 washout experiments that were recorded over a period of 4 hours.

***(A)*** Images show microtubule asters in PtK2 cytoplasts 4 hours after Nocodazole, Y27632 and Cytochalasin-D washout (top), or after Nocodazole washout, but in the presence of Y27632 (20µM), and Jasplakinolide (600nM) (bottom).
***(B)*** Time lapse sequences show the monitoring of endocytosed beads (500nm non-functionalized fluorescent polystyrene beads) over 4 hours in the various recentering conditions. In all cases, the

displacement of the centrosome is remarkably similar to the motion of the beads. Kymographs show the coordinated displacement of the centrosome and the beads inside the zoomed in region. The linescans were performed along a line following the trajectory of the centrosome and joining its initial and final positions.

***(C)*** Time lapse sequences show centrosome recentering upon all drugs washout. In this case, recentering occurs almost exclusively through the repositioning of the center of mass of the cytoplast and in the absence of absolute centrosome displacement. The dotted green line highlights the motion of the centrosome, and the dotted magenta line highlights the displacement of the center of mass.

***(B)*** Images are max projections, the centrosome is further processed with an unsharp mask and a gaussian blur.

***(C)*** Images were processed following the described pipeline in the dedicated extended material and method section.

### Supplementary Movies

#### Movie S1

Laser ablation experiment inside PtK2-GFP cells. A few microtubules are ablated on one side of the centrosome and the motion of the centrosome is recorded during the two minutes that follow ablation.
A 2 µm wide Z-stack (spaced by 1 µm) is acquired every 10 seconds and is then MAX projected onto a single plane.

#### Movie S2

Laser ablation experiment inside PtK2-GFP cytoplasts. A few microtubules are ablated on one side of the centrosome and the motion of the centrosome is recorded during the two minutes that follow ablation. A 2 µm wide Z-stack (3 steps spaced by 1 µm) is acquired every 10 seconds and is then MAX projected onto a single plane.

#### Movie S3

Repeated laser ablation experiment inside a PtK2-GFP cytoplast. Microtubules are extensively and repeatedly depleted on one side of the centrosome during five minutes and the motion of the centrosome is recorded in parallel. Actin appears in magenta and is stained using SiR Actin. A 2 µm wide Z-stack (3 steps spaced by 1 µm) is acquired every 10 seconds and is then MAX projected onto a single plane.

#### Movie S4

PtK2-GFP cell 2 hours after micro-injection with 1µM of labelled tubulin (green). Typical acquisition from which the kymographs in Figure S3A were built. The reader can visualize the interest of tubulin speckle microscopy in discriminating motion and dynamics along the lattice of the microtubules. Frames are acquired every 3 seconds over a 105 second period.

#### Movie S5

PtK2-WT cell stained using SIR-Actin (cyan), 2 hours after micro-injection of 1µM of labelled actin (magenta). Typical acquisition from which the kymographs in Figure S3B were built. The reader can visualize contraction events along stress fibers with the presence of antiparallel speckle slidings along the length of the stress fibers. Frames are acquired every 1 minute over a 30-minute period.

#### Movie S6

PtK2-WT cell 2 hours after its microinjection with both actin (magenta) and tubulin (green). Representative responses of single microtubules in the 15 seconds that followed laser ablation. On the left, a microtubule depolymerization event after laser ablation with no mechanical relaxation. On the right, actin stress fiber recoiling after laser ablation accompanied by a local microtubule buckling and recoiling. Frames are acquired every 3 seconds over a 15 second period.

#### Movie S7

PtK2-WT cell 2 hours after its microinjection with both actin (magenta) and tubulin (green). Microtubule displacement event occurring when only actin was ablated. The white “*” on the video shows the localization of the laser impact inside the actin meshwork. Frames are acquired every 3 seconds over a 15 second period.

#### Movie S8

Repeated laser ablation experiment inside a PtK2-GFP cytoplast treated with Jasplakinolide (600nm) and Y27632 (20µM) for 4 hours. Microtubules are extensively and repeatedly depleted on one side of the centrosome during five minutes and the motion of the centrosome is recorded in parallel. A 2 µm wide Z-stack (3 steps spaced by 1 µm) is acquired every 10 seconds and is then MAX projected onto a single plane.

#### Movie S9

Nocodazole washout, Y27632 washout, Cytochalasine-D washout. Representative live centrosome recentering event showing the repositioning of the centrosome and the center of mass inside a cytoplast over a time course of a few hours. 9 selected timepoints are shown from a 4 hour acquisition composed of 4 µm wide Z-stacks (5 steps spaced by 1 µm) acquired every 5 minutes and then MAX projected onto single planes. Images are processed following the described pipeline in the dedicated extended material and method section.
